# Auditory mismatch responses are differentially sensitive to changes in muscarinic acetylcholine versus dopamine receptor function

**DOI:** 10.1101/2021.03.18.435979

**Authors:** Lilian A. Weber, Sara Tomiello, Dario Schöbi, Katharina V. Wellstein, Daniel Müller, Sandra Iglesias, Klaas E. Stephan

## Abstract

The auditory mismatch negativity (MMN) has been proposed as a biomarker of NMDA receptor (NMDAR) dysfunction in schizophrenia. Pathophysiological theories suggest that such dysfunction might be partially caused by aberrant interactions of different modulatory neurotransmitters with NMDARs, which could explain heterogeneity among patients with schizophrenia and their treatment response. Understanding the differential impact of different neuromodulators on readouts of NMDAR function is therefore of high clinical relevance.

Here, we report results from two studies (*N*=81 each) which systematically tested whether the MMN is sensitive to diminishing and enhancing cholinergic vs. dopaminergic function. Both studies used a double-blind, placebo-controlled between-subject design and monitored individual drug plasma levels. Using a novel variant of the auditory oddball paradigm, we contrasted phases with stable versus volatile probabilities of tone switches. In the first study, we found that the muscarinic acetylcholine receptor antagonist biperiden reduced and/or delayed mismatch responses, particularly during stable phases of the experiment, whereas this effect was absent for amisulpride, a dopamine D2/D3 receptor antagonist. The direct comparison between biperiden and amisulpride indicated a significant drug × mismatch interaction. In the second study, neither elevating acetylcholine nor dopamine levels via administration of galantamine and levodopa, respectively, exerted significant effects on MMN.

Overall, our results indicate differential sensitivity of the MMN to changes in cholinergic (muscarinic) versus dopaminergic receptor function. This finding may prove useful for developments of future tools for predicting individual treatment responses in disorders that show abnormal MMN, such as schizophrenia.

## Introduction

The auditory mismatch negativity (MMN) is an electrophysiological response to rule violations in auditory input streams (Näätänen et al., 2001, 2011). It is commonly defined as the difference between event-related potentials (ERPs) to predictable (‘standard’) and surprising (‘deviant’) auditory events, and has been interpreted as reflecting the update of a predictive (generative) model of the acoustic environment (Winkler, 2007; Garrido et al., 2009; Lieder et al., 2013a, 2013b; Weber et al., 2020).

The auditory MMN is of major interest for translational research in psychiatry. First, there is strong evidence that MMN amplitudes are significantly reduced in patients with schizophrenia (for meta-analyses, see (Umbricht and Krljes, 2005; Erickson et al., 2016; Avissar et al., 2018)). Second, numerous studies in animals and humans have demonstrated convincingly that the MMN is sensitive to pharmacological alterations of NMDA receptor (NMDAR) function (Javitt et al., 1996; Umbricht et al., 2000; Heekeren et al., 2008; Schmidt et al., 2012; Rosburg and Kreitschmann-Andermahr, 2016) – which, in turn, plays a major role in pathophysiological theories of schizophrenia (Olney and Farber, 1995; Friston, 1998; Goff and Coyle, 2001; Stephan et al., 2006, 2009; Corlett et al., 2011, 2016; Javitt, 2012; Friston et al., 2016). The MMN has thus been suggested as a potential readout of NMDA receptor (NMDAR) hypofunction in schizophrenia and has been proposed as a promising translational biomarker (Light and Näätänen, 2013; Todd et al., 2013; Näätänen et al., 2015).

Here, we investigate whether the auditory MMN is differentially sensitive to cholinergic versus dopaminergic challenges. Acetylcholine (ACh) and dopamine (DA) are two modulatory transmitters with a general capacity to modulate NMDAR function ((Hallett et al., 2006; Lin et al., 2010; Zappettini et al., 2014; Zwart et al., 2018) for review, see (Gu, 2002)), and their relative contribution to NMDAR *dys*regulation has been suggested as a major cause of heterogeneity in clinical trajectories among patients with schizophrenia (‘dysconnection hypothesis’, (Stephan et al., 2006, 2009).

Understanding the substantial heterogeneity within patient populations under the current syndromatic diagnostic categories is one of the main challenges for psychiatry and an essential basis for individualized treatment predictions (Kapur et al., 2012; Stephan et al., 2017). As a consequence, biomarkers are sought that *differentiate* between alternative pathophysiological mechanisms where, ideally, these mechanisms relate to different available treatment options.

Critically, detecting alterations of cholinergic and dopaminergic neuromodulatory transmitter systems may indeed be relevant for treatment choice in schizophrenia: while standard antipsychotic treatment options in schizophrenia rely on antagonism at D2/D3 dopaminergic receptors, they show considerable variability in their binding capacity to other receptors (Nasrallah, 2008). Most notably, some of the most potent antipsychotics (olanzapine and clozapine) have strong affinity to cholinergic (specifically: muscarinic) receptors (Lavalaye et al., 2001), in contrast to almost all other second generation antipsychotics. Therefore, a readout of the functional status of muscarinic vs. dopaminergic systems in the individual could prove valuable for understanding the neurobiological basis of differential treatment responses in schizophrenia, and, subsequently, for guiding treatment (Stephan et al., 2009, 2015).

However, whether such a readout of muscarinic vs. dopaminergic function could be obtained from MMN responses is not clear. While nicotinic stimulation has been demonstrated to enhance MMN amplitudes (Harkrider and Hedrick, 2005; Inami et al., 2005, 2007; Baldeweg et al., 2006; Dunbar et al., 2007; Martin et al., 2009; Fisher et al., 2012; Knott et al., 2012; Hamilton et al., 2018), the role of muscarinic cholinergic receptors for MMN is less well established. The few human studies investigating the effects of muscarinic antagonists scopolamine and biperiden on auditory mismatch processing were inconclusive and showed mixed results (Pekkonen et al., 2001, 2005; Klinkenberg et al., 2013; Caldenhove et al., 2017). Similarly, while several pharmacological studies of DA failed to show significant effects on MMN (Kähkönen et al., 2002; Leung et al., 2007, 2010; Korostenskaja et al., 2008), other studies reported significant alterations of MMN by antipsychotic drug treatment, hinting at a possible effect of DA (Kähkönen et al., 2001; Zhou et al., 2013). However, the latter interpretation is vague, given that the antipsychotic drugs studied affect numerous types of receptors.

In summary, there is inconclusive evidence concerning the sensitivity of the auditory MMN to dopaminergic and muscarinic alterations. This could be due to small sample sizes, unspecific drugs (such as antipsychotics), and/or individual differences in pharmacokinetics and thus variability in actual drug plasma levels across participants.

Here, we report results from two double-blind, between-subject, placebo-controlled studies that address these problems and test whether the MMN is differentially sensitive to cholinergic and dopaminergic alterations. In study 1 (N=81), we tested the effects of biperiden, a selective muscarinic M1 receptor antagonist, on mismatch related ERPs, and compare them to the effects of amisulpride, a selective dopaminergic D2/3 receptor antagonist. In study 2, we employed exactly the same study design, paradigm, and analysis strategy in a separate sample (N=81), to test the impact of elevated cholinergic vs. dopaminergic transmission on MMN amplitudes, contrasting the acetylcholinesterase inhibitor galantamine to the dopamine precursor levodopa. In both studies, we used estimates of the actual drug plasma levels at the time participants performed the experimental task in order to account for individual differences in pharmacokinetics.

Importantly, we used a new variant of an auditory oddball paradigm with explicitly varying levels of stability over time. The MMN has previously been found to be sensitive to the overall level of volatility within a block, such that MMN amplitudes were higher in blocks with more stable regularities (Todd et al., 2014; Dzafic et al., 2020). We were interested in whether cholinergic manipulations would interact with this sensitivity to environmental volatility. This was motivated by theoretical accounts (Mathys et al., 2011) and experimental findings that volatility affects precision-weighting of prediction error responses, possibly via interactions of ACh and NMDA receptor dependent mechanisms (Iglesias et al., 2013; Weber et al., 2020). While these latter studies employed trial-by-trial computational models, our paradigm was designed to contain distinct phases of volatility. This allowed us to capture the interaction between environmental volatility and mismatch processing using the conventional approach of trial averaging and comparing mismatch responses across different phases of volatility, which maximizes sensitivity for detecting volatility effects. Here, we focus on the results of this conventional analysis, but also present the (complementary) model-based perspective on our data in the supplementary material.

In brief, our results suggest that muscarinic receptors play a critical role for the generation of MMN responses and their dependence on environmental volatility, whereas no such evidence was found for dopamine receptors.

## Methods: Study 1

### Participants

In total, 81 volunteers (mean age 22.7 years (SD=3.6, range=18-38)) participated in study 1. In this initial study with its focus on the feasibility of an EEG-based readout of differential sensitivity to cholinergic (muscarinic) vs. dopaminergic function, we aimed for controlling potential confounds as tightly as possible. In addition to measuring individual drug plasma levels and transmitter-relevant single nucleotide polymorphisms (see below), we therefore only recruited male participants in order to avoid the significant influence of fluctuating estrogen levels on dopaminergic and cholinergic systems (Gasbarri et al., 2012; Colzato and Hommel, 2014; Barth et al., 2015). However, this has the obvious disadvantage that our study is not representative for the entire population. This is a significant limitation which we revisit in the Discussion. All participants were right-handed, Caucasian, and non-smokers with normal or corrected-to-normal vision. Further exclusion criteria included serious chronic or current physical or mental illness, drug consumption, and hearing aids.

To exclude any cardiac abnormalities that could render a pharmacological intervention risky, participants underwent a clinical examination including electrocardiogram (ECG) before data acquisition. Participants were randomly assigned to one of three drug groups: placebo, amisulpride, or biperiden (between-subject design, N=27 per group), with both the participant and the experimenters blind to the drug label. All participants gave written informed consent prior to data acquisition and were financially reimbursed for their participation. The study was approved by the cantonal Ethics Committee of Zurich (KEK-ZH-Nr. 2011-0101/3).

Data from a total of ten participants could not be used in the group analysis presented here for the following reasons: change of the stimulus sequence after the first few participants (N=6), technical issues during measurement (N=2), and failure to sufficiently correct for eye blink artefacts during preprocessing of EEG data (N=2, see below). Therefore, the results reported here are based on a final sample of N=71 participants, with N=25 in the placebo group (mean age 23.2 years (SD=4.8, range=18-38)), N=24 in the amisulpride group (mean age 22.4 years (SD=3.4, range=18-33)), and N=22 in the biperiden group (mean age 22.5 years (SD=3.1, range=18-29)). Criteria for excluding data sets from the group analysis were defined and documented in a time-stamped analysis plan prior to un-blinding of the analyzing researcher (see below, section ‘Analysis Plan, Data and Code Availability’).

### Pharmacological Substances & Administration

At the clinical examination, participants were instructed to abstain from the consumption of alcohol and grapefruit juice for 24h before the EEG measurement, not to take any medications within 3 days before the experiment and not to consume other drugs. They were further instructed not to eat for 3h before the EEG measurement, and to abstain from driving a car for 48h after the experiment.

Approximately 80min before the start of the EEG measurement, capsules of each compound (amisulpride/biperiden/placebo) were administered as a single oral dose. All capsules had the same visual appearance and drug administration was conducted in a double-blind fashion. The drugs were prepared by the local pharmacy Bellevue Apotheke, Zurich.

Amisulpride was administered using Solian® 400mg mixed with 570mg of lactose. At this dose, amisulpride blocks postsynaptic D_2_ and D_3_ receptors, thus inhibiting DA transmission (Chhabra and Bhatia, 2007). Biperiden capsules contained two units of 2mg Akineton® (i.e., 4 mg in total) mixed with 880mg of lactose. Biperiden is the most selective M1 antagonist available for human subjects (Katayama et al., 1990; Bolden et al., 1992) and has only minor peripheral anticholinergic effects in comparison with other anticholinergic substances. Placebo capsules contained 960mg of lactose.

### Blood samples

Four blood samples were collected per participant in order to (1) estimate the actual drug plasma levels at the time participants performed the experimental task (using two samples), and (2) to assess genetic variation at functional single nucleotide polymorphisms (SNPs) of two genes relevant to the pharmacological intervention (using two samples). However, the assessment of genetic effects in our study is constrained by the very limited sample size. In particular, for some genotypes of interest, there were only 2 or 3 individuals within certain drug groups showing these genotypes. We therefore refrain from interpreting or discussing these genetic effects any further and report them in the supplementary material (section S2.3) for completeness and potential guidance for future follow-up studies with larger sample sizes.

#### Drug plasma concentration

For both pharmacological agents, the expected maximal plasma concentration was around 1h after intake (amisulpride: first peak of plasma concentration after 1h, second peak at 3-4h, absolute bioavailability of 48%, elimination half-life ~12h (https://compendium.ch/mpro/mnr/8962/html/de); biperiden: for single dose usage, peak of plasma concentration around 1h after administration, absolute bioavailability ~33%; elimination half-life 11-21.3h (https://compendium.ch/mpro/mnr/1853/html/de)).

The first blood sample was collected on average 75.67min (SD=3.22) after drug intake. A second blood sample was taken on average 188.99min (SD=9.91) after drug administration. Blood samples were collected in tubes containing heparin as anticoagulant, centrifuged at 10°C for 10min at 3000xg and finally stored at −86°C until analysis.

Blood analysis was performed by the Institute of Clinical Chemistry at the University Hospital Zurich with a detection threshold of 1nmol/L. Samples were measured using liquid chromatography coupled to tandem mass spectrometry (LC-MS/MS). Methods were fully validated and accredited according to ISO 17025. The lower limits of quantification were for amisulpride 2 nmol/L, for biperiden 1 nmol/L, for galantamine 1 μg/L, and for levodopa 10 μg/L.

Estimated drug plasma levels at the time of the experimental task were read off a linear approximation of drug concentration decay between the two collection time points for each individual and entered the group level general linear model (GLM) as a covariate (see below).

### Paradigm

Participants passively listened to a sequence of tones, presented binaurally through headphones, while engaging in a visual distraction task (described below). The auditory stimuli consisted of two pure sinusoidal tones; a high (528Hz) and a low (440Hz) tone. A total of 1800 tones were presented, with a duration of 70ms each and an inter-stimulus interval of 500ms, see Figure 1 for a visualization of the paradigm and relative timing of events. Auditory and visual stimuli were presented using PsychToolbox (PTB3, psychotoolbox.org).

**Figure 1.**
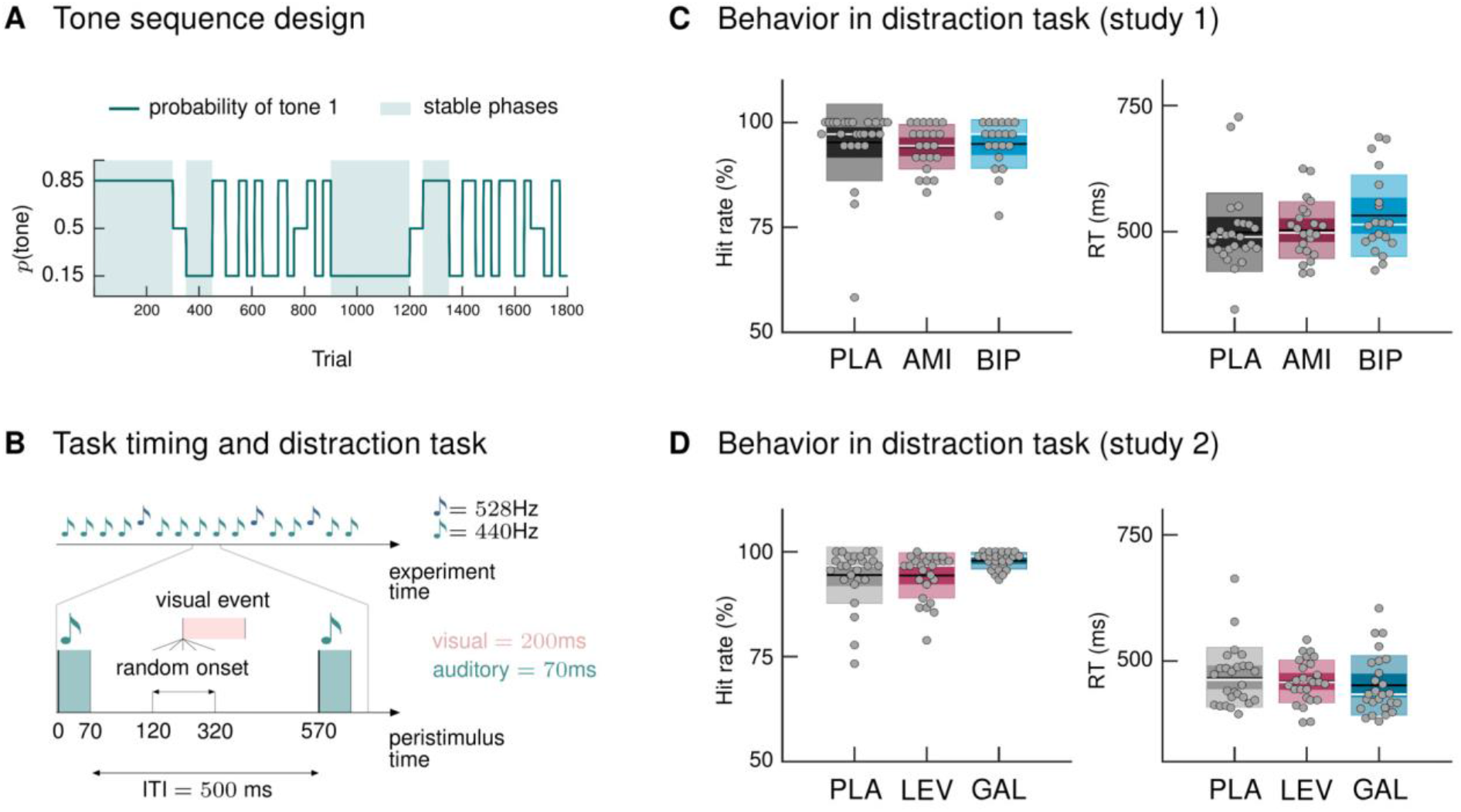
Paradigm and behavioral results. **A** Probability structure for the tone sequence in the oddball MMN paradigm with volatility. The probability of hearing the higher tone (tone 1, with *p*(tone 2) = 1 − *p*(tone 1)) varied over the course of the tone sequence as indicated by the blue line. Tone 1 functioned as the ‹deviant› in phases where it was less likely (*p* = 0.15), and as the ‹standard› when it was more likely than tone 2 (*p* = 0.85). Stable phases (*p* constant for 100 or more trials) alternated with volatile phases (*p* changes every 25-60 trials). **B** Experimental task: Overview of timing of events. Participants passively listened to a sequence of 1800 tones while performing a visual distraction task. Visual events occurred after tone presentations at a randomly varying delay between 50 and 250ms after tone offset, in 36 (study 1) and 90 (study 2) out of 1800 trials. ITI = Inter-stimulus interval. **C, D** Hit rates and reaction times, per drug group, for the visual distraction task, plotted using the notBoxPlot function (https://github.com/raacampbell/notBoxPlot/). Mean values are marked by black lines; medians by white lines. The dark box around the mean reflects the 95% confidence interval around the mean, and the light outer box 1 standard deviation. **C** There were no significant differences between drug groups in performance on the visual distraction task. **D** Participants in the galantamine group had higher hit rates in the distraction task (see main text). PLA = placebo, AMI = amisulpride, BIP = biperiden, LEV = levodopa, GAL = galantamine group.

#### Auditory oddball sequence

We used a new variant of the auditory oddball paradigm, in which we explicitly varied the degree of volatility in the auditory stream over time. In a classical oddball paradigm, one stimulus is less likely to occur and thus considered a surprising, or ‘deviant’, stimulus, whereas the other stimulus is considered the ‘standard’ event. Our sequence was generated such that both tones could be perceived as standard (predictable) or deviant (surprising), depending on the current context. More specifically, the probability of hearing the high tone was either 0.15 (in which case it was the deviant) or 0.85 (in which case it functioned as a standard), except for four short phases, 50 trials each, in which the probability of hearing either tone was equal. Critically, the tone sequence comprised ‘stable’ phases in which probabilities remained constant over 100 or more trials, and ‘volatile’ phases in which the probability changed every 25-60 trials. Figure 1A displays the probability structure underlying the tone sequence and the division into stable and volatile phases.

In order to ensure that both tones appeared equally often in both roles, the second half of the stimulus stream was a repetition of the first half, with only the tones switched. This avoids potential confounding effects by ensuring that both stimulus categories have, on average, the same physical properties across the duration of the experiment.

#### Visual distraction task and behavioral data analysis

Participants performed a distracting visual task and were instructed to ignore the sounds, following the suggestion that MMN assessment is optimal when the participant’s attention is directed away from the auditory domain (Näätänen, 2000). The task consisted of detecting changes to a centrally presented small white square. Whenever the square opened to either the left or the right side, participants were instructed to press a button on a response box with their index finger (left opening) or middle finger (right opening). There were 36 ‘square openings’ in total (half of them to the left), occurring at irregular intervals. They did not coincide with tone presentations but always followed a tone with a delay varying randomly between 50 and 250ms after tone offset (see Figure 1B).

Based on the participants’ responses, we calculated mean reaction times and hit rates (defined as the proportion of correct responses relative to the total number of visual targets). Due to technical issues during measurement, behavioral data from three participants (N=1 amisulpride group, N=2 biperiden group) were missing. We cannot exclude the possibility that these participants, as well as one participant with very low performance level on the distraction task (hit rate<75%, N=1, placebo group, see Figure 1C), were paying attention to the auditory input instead of focusing on the visual task. Nevertheless, we decided to include these data sets in the group level analysis for the following reasons: (i) Effects of attention on MMN are generally small and not always consistent (Näätänen, 2000; Chennu et al., 2013; Auksztulewicz and Friston, 2015; Hsu et al., 2015; Garrido et al., 2017), (ii) it is unlikely that these N=4 datasets are driving the effects (or absence thereof) observed in the whole sample (N=71), and (iii) distraction tasks employed in auditory mismatch paradigms vary considerably. For most of them, including the one used here, it cannot be excluded that participants do attend to the tones occasionally, even when otherwise engaging in the distraction task properly. However, for transparency, we report the results of our group analysis when excluding the four data sets in the supplementary material (section S2.2).

### EEG data acquisition and preprocessing

EEG data were collected at a sampling rate of 500Hz using an EASYCAP system 64 scalp electrodes including one electrooculography (EOG) channel (10-20 layout; EASYCAP GmbH, https://www.easycap.de/wordpress/). Data were recorded with nose-reference. Before starting the experimental task, impedances were ensured to be well below 20kOhm for all channels. For a subset of participants, ECG and pulse oximetry data were additionally acquired via a bipolar amplifier (BrainAmp ExG; Brain Products GmbH, https://www.brainproducts.com/index.php), however, these data were not analyzed in the present study. For one participant, erroneous cabling during data acquisition resulted in a different order of EEG channels. This could be corrected for during the pre-processing of the data.

Pre-processing and data analysis of EEG data were performed using SPM12 (v6906, http://www.fil.ion.ucl.ac.uk/spm/) and Matlab (R2018b). Continuous EEG recordings were re-referenced to the average, high-pass filtered using a Butterworth filter with cutoff frequency 0.1Hz, down-sampled to 250Hz, and low-pass filtered using Butterworth filter with cutoff frequency of 30Hz.

The data were epoched into 500ms segments around tone onsets, using a pre-stimulus time window of 100ms. We did not baseline-correct the epochs. Whether the benefits of traditional baseline correction outweigh its downsides is still a matter of debate (Alday, 2019). Here, we wanted to avoid mixing anticipation or prediction signals with event-related responses, which we interpret as learning or model update signals. However, to explore the robustness of our findings under different choices of analysis strategy, we also present results under an alternative pre-processing pipeline which included strong high-pass filtering and baseline correction (supplementary material section 2.4)

A vertical EOG channel was computed as the difference between channel Fp1 and the EOG channel which was placed beneath the left eye. We accounted for eye-movement related artefacts by applying the signal space projection (SSP) eye blink correction method (Nolte and Hämäläinen, 2001) as implemented in SPM12: This approach uses an estimate of the spatial topography due to eye activity to define ocular source components and removes eye activity by regressing these components out of the EEG data.

In particular, eye blink events were identified with a thresholding approach applied to the data from the vertical EOG channel. Detected eye blink events were used to epoch the continuous EEG into 1000ms segments around these events, excluding any epochs containing large transients. Ocular components were determined using singular value decomposition (SVD) of topographies from all the eye blink trials and all the time points. The leading SVD component was used to define the noise subspace that was subsequently projected out of the data (Nolte and Hämäläinen, 2001). This projection was applied to the data epoched around the auditory stimulus presentation. For all participants, we verified that the leading SVD component had the typical spatial topography of an eye blink artifact and resulted in satisfactory eye blink correction performance (inspected visually by plotting the average eye blink in a subset of channels before correction and after correction). To achieve this, in a subset of participants, the default eye blink detection threshold of 5 SD was changed to a value that resulted in improved correction performance. Participants for which such a component could not be identified were excluded from further analysis (N=2: one in the amisulpride group, one in the biperiden group).

Finally, epochs in which the (absolute) signal recorded at any of the channels exceeded 75μV were removed from subsequent analysis. For all channels in all participants, the number of excluded epochs was below 20% of the total number of epochs. The number of remaining good trials was 1775 on average across participants (SD=28) and almost identical across drug conditions (placebo group: 1775, SD=29; amisulpride group: 1775, SD=27; biperiden group: 1776, SD=30).

The remaining good trials were converted, for each participant, into scalp images for all 63 EEG channels and all time points between 100ms and 400ms after tone onset, using a voxel size of 4.2mm × 5.4mm × 4.0ms. The images were spatially smoothed with a Gaussian kernel (FWHM: 16mm × 16mm) in accordance with the assumptions of Random Field Theory (Worsley et al., 1996; Kiebel and Friston, 2004) and to accommodate for between-subject spatial variability in channel space.

### First level general linear model

We defined categorical trial types based on our tone sequence: deviant trials (defined as the first tone with a different frequency; following previous studies (Garrido et al., 2008), we only considered deviants presented after at least 5 repetitions (N=119) and standard trials were defined as the 6th repetition of the same tone, to keep trial numbers comparable across conditions (N=106). Based on the probability structure of the input sequence, we further divided these into deviants into stable phases, deviants in volatile phases, standards in stable phases, and standards in volatile phases. Stable phases were defined as phases in which the probability of hearing the high tone did not change for at least 100 trials; volatile phases were all other phases of the experiment. Note that while we chose this trial definition due to its specificity, we also considered alternative trial definitions (specified a priori as part of our analysis plan) which allow for more trials per condition to be retained in the EEG analysis. In the supplementary material (section S2.4), we present results under such an alternative definition.

Per participant, we modeled the trial-wise 3D ERP images with a GLM which implements a factorial design with two factors: ‘Mismatch’ (levels: 1. standards, 2. deviants) and ‘Stability’ (levels: 1. stable, 2. volatile). With regard to non-sphericity correction at this single-subject level, we assumed that the error might have different variance (i.e., non-identity) but is not correlated (independence) across conditions, in line with the recommendations in the SPM manual (http://www.fil.ion.ucl.ac.uk/spm/). This GLM only served to provide the contrast images to be used in the group level GLM. We computed contrast images (using *t*-tests) for the following contrasts of interest:

- mismatch effect: standards vs. deviants
- stability effect: stable vs. volatile
- interaction effect: stable mismatch vs. volatile mismatch
- stable mismatch: stable standards vs. stable deviants
- volatile mismatch: volatile standards vs. volatile deviants

For visualization purposes, grand average waveforms were computed for each condition.

Additionally, and in line with our analysis plan, we performed a model-based analysis in which the conventional trial definition (‘standard’ versus ‘deviant’ trials) was replaced with a trial-wise estimate of the amount of prediction error that each tone in our sequence would elicit, according to an ideal observer model. This analysis has the advantage of taking into account the trial-by-trial dynamics; on the other hand, it requires making assumptions about the nature of the learning process. The details of this analysis and the obtained results are presented in the supplementary material (sections 1.4 and 2.4).

### Group level general linear models

Random effects group analysis across all participants was performed using a standard summary statistics approach (Penny and Holmes, 2007). We used a separate group-level GLM for each effect of interest from the first level GLM, which implements a factorial design with the between-subject factor ‘drug’ (levels: 1. placebo, 2. amisulpride, 3. biperiden). With regard to non-sphericity correction, the group-level analysis assumed independence (measurements are unrelated to each other), given the between-subject design, and non-identity (variances may differ across measurements).

We introduced a covariate for the estimated drug plasma concentration levels of both pharmacological agents, where we allowed for an interaction with the drug factor and mean-centered the covariate within drug groups (for the reasoning behind this, please refer to section S1.1 in the supplementary material).

In sum, our design effectively comprised two within-subject factors – mismatch (standards vs. deviants) and stability (stable vs. volatile), which we specified in our first-level GLM – and one between-subject factor, drug group (placebo vs. amisulpride vs. biperiden). At the group level, we were particularly interested in the interaction between the factors mismatch and drug, and the three-way interaction between mismatch, stability, and drug.

#### Pharmacological effects

For each effect of interest from the first level, we used 8 separate *t*-tests to examine: average positive and negative EEG deflections for the effect across drug groups, and drug differences in the expression of the effect: amisulpride compared to placebo in both directions, biperiden compared to placebo in both directions, and differences between amisulpride and biperiden in both directions.

In addition to an initial analysis across the whole time-sensor space, we investigated drug effects within a smaller, functionally constrained search volume, which comprised those regions of the time × sensor space where we found significant average effects (across drugs). Specifically, a mask was functionally defined for each effect of interest and created by combining the images of significant activations for the positive and the negative average effect (logical OR) of that contrast. Importantly, the differential contrasts used to test for drug effects were orthogonal to the average contrasts used to construct these masks.

For all analyses, we report all results that survived family-wise error (FWE) correction, based on Gaussian random field theory, across the entire volume (time × sensor space), or within the functional masks (small volume correction, SVC), at the peak level (p<0.05).

### Analysis plan, data and code availability

Prior to the unblinding of the researcher conducting the analysis, a version-controlled and time-stamped analysis plan was created. This plan detailed the analysis pipeline ex ante (see next sections). The analysis plan is provided online at https://gitlab.ethz.ch/tnu/analysis-plans/weber-muscarinic-mmn-erp. The data used for this manuscript are available at https://research-collection.ethz.ch/handle/20.500.11850/477685, adhering to the FAIR (Findable, Accessible, Interoperable, and Re-usable) data principles. Furthermore, the analysis code that reproduces the results presented here is publicly available on the GIT repository of ETH Zurich at https://gitlab.ethz.ch/tnu/code/weber-muscarinic-mmn-erp-2021. The code used for running the experimental paradigm will also be made publicly available, as part of a future release of the open source software package TAPAS (www.translationalneuromodeling.org/tapas).

## Methods: Study 2

Study 2 employed exactly the same study design as study 1 except for the pharmacological agents used. The participants did not overlap across studies. In the following, we only report the parts of the experiment that differed to study 1 and refer the reader to study 1 for all other aspects of the experiment and analysis. In particular, we followed exactly the same analysis steps as outlined in the analysis plan for study 1 (see section ‘Analysis plan, data and code availability’).

### Participants

In total, 81 male volunteers (mean age 23.5 years (SD=3.5, range=18-35)) participated in study 2. Inclusion and exclusion criteria were identical to study 1.

Participants were randomly assigned to one of three drug groups: placebo, levodopa, or galantamine (between-subject design, N=27 per drug group) with both participants and experimenters blind to the drug label. All participants gave written informed consent prior to data acquisition and were financially reimbursed for their participation. The study was approved by the cantonal Ethics Committee of Zurich (KEK-ZH-Nr. 2011-0101/3).

Data from three participants could not be used in the group analysis presented here due to a diagnosis of diabetes (N=1; prior to unblinding, we decided not to analyze this dataset because of potential interactions of insulin with DA; (Figlewicz et al., 2003; Fiory et al., 2019), technical issues during measurement (N=1), and an adverse event prior to data acquisition (N=1; nausea). Therefore, the results reported here are based on a sample of N=78 participants, with N=26 in the placebo group (mean age 24.3 years (SD=3.9, range=19-35)), N=26 in the levodopa group (mean age 23.6 years (SD=3.8, range=19-33)), and N=26 in the galantamine group (mean age 22.7 years (SD=3.0, range=18-33)).

### Pharmacological substances, Administration & Blood samples

Approximately 80min before the start of the EEG measurement, capsules of each compound (levodopa/galantamine/placebo) were administered as a single oral dose. All capsules had the same visual appearance and drug administration was double-blind.

For levodopa, we followed closely the procedure reported by (Rihet et al., 2002) by using a single oral dose administration of Madopar® DR (Roche Pharma (Switzerland) AG, 4153 Reinach; Licence number: 53493 (Swissmedic)), mixed with 670mg lactose. Madopar DR is a dual-release formulation containing 200mg levodopa and 50mg benserazide. Levodopa is the immediate metabolic precursor of DA and is decarboxylated to DA both in the central (CNS) and the peripheral nervous system. Concurrent administration of benserazide, a dopa decarboxylase inhibitor, which does not cross the blood-brain barrier, reduces the extracerebral side effects of levodopa and enhances the amount of levodopa reaching the CNS (Crevoisier et al., 1987). Galantamine was administered as a single oral dose of Reminyl® (Janssen-Cilag (Switzerland) AG, Baar, ZG; Licence number: 56754 (Swissmedic)) containing 8mg of galantamine, mixed with 920mg lactose. As a selective, competitive and reversible inhibitor of acetylcholinesterase (AChE), an enzyme which degrades ACh, galantamine increases the availability of ACh. Additionally, it may act as a positive allosteric modulator of nicotinic receptors (Schrattenholz et al., 1996; Samochocki et al., 2003) although this property is being debated (Kowal et al., 2018). Placebo capsules only contained lactose. Drugs were prepared by the local pharmacy Bellevue Apotheke, Zurich.

For both pharmacological agents, the expected maximal plasma concentration was around 1h after intake (levodopa: peak of plasma concentration after 1h, absolute bioavailability of ~78% when using the dual-release formulation, elimination half-life ~1.5h (https://compendium.ch/product/56931-madopar-dr-tabl-250-mg/mpro); galantamine: peak of plasma concentration around 1-2h after administration, absolute bioavailability ~88.5%; elimination half-life 7-8h (https://compendium.ch/product/1018816-reminyl-prolonged-release-kaps-8-mg/MPro)).

The first blood sample was collected on average 77.71 min (SD: 14.38) after drug intake. A second blood sample was taken 192.79 min (SD: 18.45) after drug administration. Blood samples were collected and processed as described in study 1. As in study 1, an additional blood sample was collected for assessing genetic variation at selected functional single nucleotide polymorphisms (SNPs). As for study 1, we report the results of the genetic analyses in the supplementary material (section S2.3), bearing in mind the sample size limitations mentioned above.

### Paradigm

We used the same paradigm and distraction task as in study 1. However, following observations during study 1 that participants found the task rather tiring due to long sequences without visual events, we increased the number of square openings in the visual distraction task from 36 to 90 to make the task more engaging. One participant had a hit rate below 75% (see section Results). Again, we report the group level results including data from this participant in the main text, but also report the results based on the analysis without this dataset in the supplementary material (section S2.2).

### EEG recording and statistical analysis

EEG recording setup, preprocessing pipeline and statistical analysis were identical to study 1. For all channels in all participants, the number of excluded epochs was below 20% of the total number of epochs, therefore, we did not mark any channels as bad. The number of remaining good trials was 1753 on average (SD=71), with no significant differences (one-way ANOVA F=1.18, p=0.31) across groups (placebo: 1770, SD=51; levodopa: 1740, SD=86; galantamine: 1750, SD=71). The specification of first level and group level GLMs was identical to study 1.

## Results: Study 1

### Behavior in distraction task

Participants reacted to visual targets on average after 509.8ms (SD=72.6) and responded correctly to 94.8% (SD=7.0) of the presented targets. There were no significant differences between drug groups in their performance on the distraction task, as assessed with a one-way ANOVAs for reaction times (*F*=1.32, *p*=0.27) and a Kruskal Wallis test for hit rates (χ^2^=2.92, *p*=0.23, Figure 1C).

### ERP effects

In the following, we report the group-level effects for our experimental factors mismatch, stability (both within-subject) and drug (between-subject), and their interactions. Here, we focus on the main effect of mismatch, the interaction of mismatch with drug, the two-way interaction mismatch × stability, and the three-way interaction mismatch × stability × drug. In the supplementary material, we additionally report the main effect of stability and its interaction with drug.

#### Main effect of mismatch

Averaging across drug groups, the strongest effect of mismatch corresponded to the classical mismatch negativity: in a large cluster of frontal, fronto-central, and central sensors, ERPs to standard tones were significantly more positive than ERPs to deviant tones from 100ms to 232ms after tone onset, with a peak difference at 172ms (*t*=16.59, *p*<0.001). The reverse was true at pre-frontal (100ms to 236ms, peak at 168ms, *t*=13.93, *p*<0.001) and temporo-parietal sensors (100ms to 328ms, peak at 176ms, *t*=14.42, *p*<0.001). We found eight additional clusters of significant differences between standard and deviant ERPs at later time points within peristimulus time, which are listed in Table 1 and partly displayed in Figure 2.

**Table 1.**
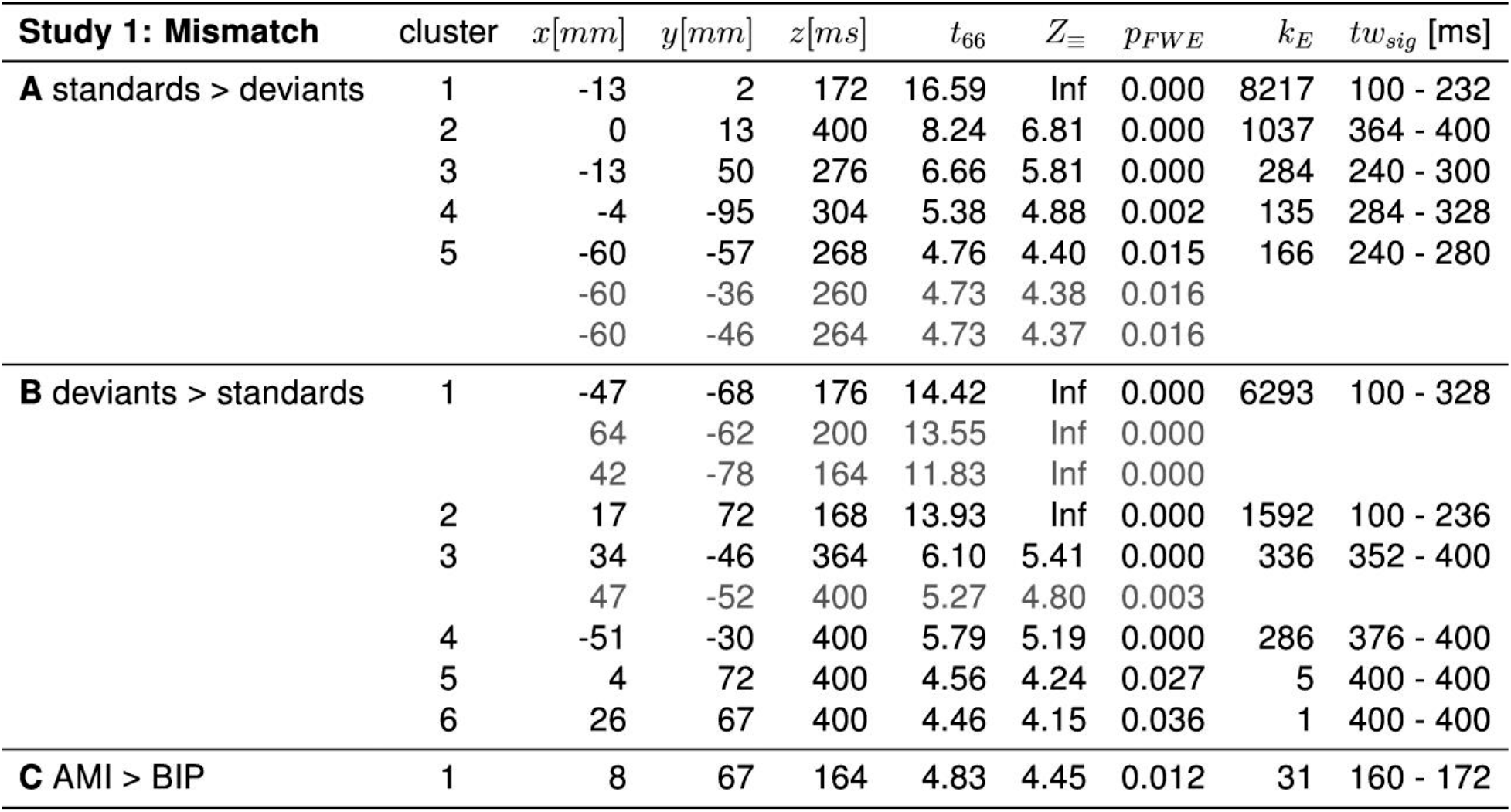
Significant clusters of activation for effects of mismatch (standards versus deviants) and pharmacological effects on mismatch in study 1. The table lists the peak coordinates (*x*, *y*, and *t* for time), peak *t* values, corresponding *Z* values, whole-volume FWE-corrected *p*-values at the peak level, and cluster size (*k_E_*). The last column lists the minimal and maximal time points of the cluster, i.e., the significant time window *t*_sig_.

**Figure 2.**
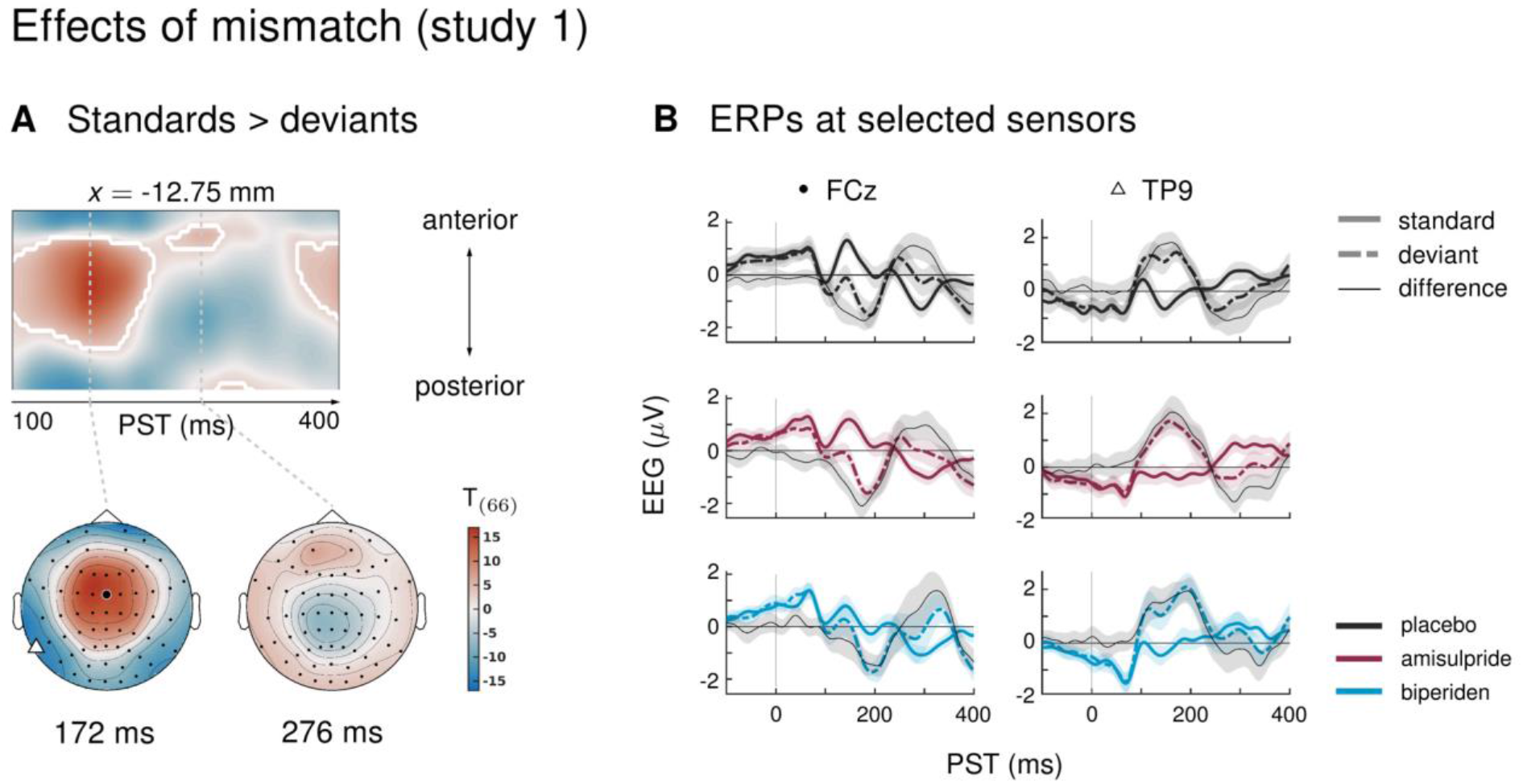
Main effect of mismatch in study 1. **A** Regions of the time × sensor space where ERPs to standards were more positive than ERPs to deviants. Displayed are *t*-maps for the contrast standards > deviants. The first map runs across the scalp dimension *y* (from posterior to anterior, y-axis), and across peristimulus time (x-axis), at the spatial x-location indicated above the map. Significant *t* values (*p* < 0.05, whole-volume FWE-corrected at the peak-level) are marked by white contours. The scalp maps below show the *t*-map at the indicated peristimulus time point, corresponding to the peak of that cluster, across a 2D representation of the sensor layout. ERPs to deviants were significantly different from ERPs to standards in large parts of the time × sensor space, including the classical mismatch negativity in fronto-central channels between 100 and 250 ms after tone onset. **B** ERPs and difference waves for selected sensors, separately for the three drug groups. The location of the chosen sensors on the scalp is marked on the scalp map in panel A by the corresponding symbol.

#### Interaction mismatch × drug

Mismatch effects were different between drug groups: biperiden delayed and topographically shifted mismatch signals compared to amisulpride and placebo (see Figure 3C for selected sensors, and Figure 3D for selected time points). When considering the whole time × sensor space and correcting for multiple comparisons using Gaussian random field (GRF) theory, this difference was significant at pre-frontal sensors for the comparison between the amisulpride and the biperiden group: between 160ms and 172ms after tone onset, the difference between standard and deviant ERPs was significantly smaller in the biperiden group compared to the amisulpride group, peaking at 164ms (*t*=4.45, p=0.012, Figure 3A, Table 1).

**Figure 3.**
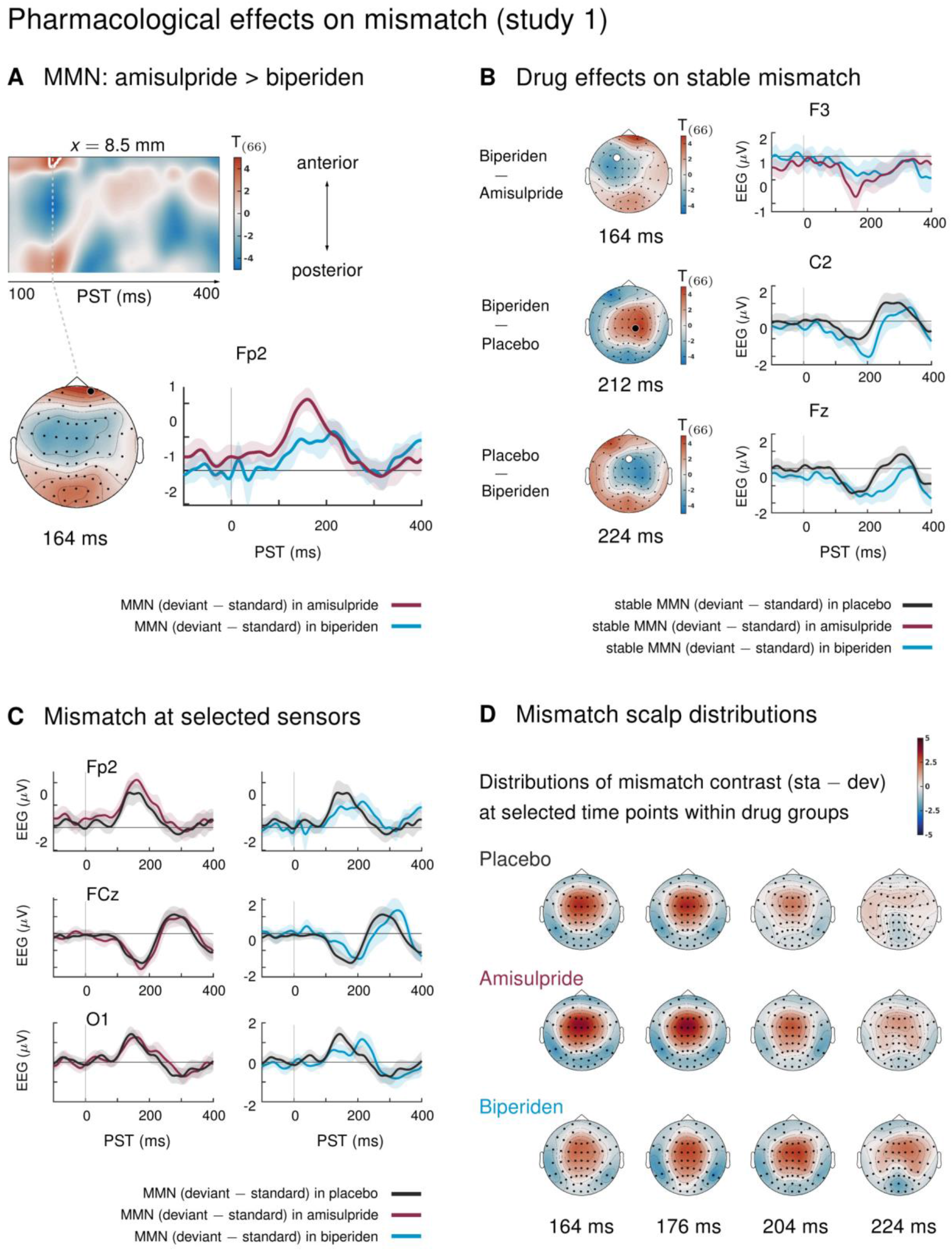
Pharmacological effects on mismatch ERPs in study 1. Logic of display as in Figure 2. **A** Mismatch response in pre-frontal sensors were significantly weaker in the biperiden group compared to the amisulpride group. Panel **B** shows additional pharmacological effects when only testing mismatch ERPs during stable phases. Panels **C** and **D** further visualize the altered mismatch response in the biperiden group (averaging across stable and volatile phases): Panel **C** displays difference waves (deviants – standards) at selected sensors for the different drug groups; Panel **D** plots the scalp distribution of the mismatch contrast at selected time points. The mismatch response in the biperiden group peaked later and more towards right central channels than in the other groups.

This did not change when constraining the search volume to the significant average mismatch effect using the functionally defined mask. However, because mismatch effects in our large sample were significant in large portions of the time × sensor space, this mask was rather unspecific. We therefore decided to deviate from our a priori analysis plan and constrain our search volume further by considering only those parts of the time × sensor space which both showed significant effects of mismatch in our sample *and* corresponded to the classical time windows and sensor locations for the mismatch negativity. In particular, we used the large cluster of frontal, fronto-central and central sensors described above which showed significant mismatch negativity between 100ms and 232ms (peak *t*=16.59) as a mask to constrain the search volume and subsequently constrain the multiple comparison correction to this volume using SPM’s small volume correction (SVC). When focusing on this subspace, an additional cluster showed a significant effect of drug on mismatch: mismatch signals were stronger in the biperiden group compared to the placebo group at right central and centro-parietal sensors with peak difference at 200ms (*t*=3.72, *p*=0.048 after SVC). This difference is indicative of both a delay and a shift in topography of mismatch signals in the biperiden group compared to the other two groups, leading to weaker mismatch (reduced MMN) early on, particularly in pre-frontal and frontal channels, but stronger mismatch (larger MMN) later on, particularly in right centro-parietal channels (see Figure 3D for a visualization).

#### Interaction mismatch × stability

We found significant interaction effects, in other words, the amount of mismatch depended on the current level of stability in the sensory input, in 3 clusters. Between 180ms and 220ms after tone onset, mismatch was significantly stronger in stable as compared to volatile phases, with a peak difference at 204ms (*t*=5.13, *p*=0.001) at central and centro-parietal sensors. Right parietal and left temporo-parietal sensors, which generally show the mismatch effect with the opposite sign compared to fronto-central channels, also showed stronger (negative) mismatch for stable phases than for volatile phases (right parietal cluster: 188-236ms, peak at 200ms, *t*=5.07, *p*=0.001; left temporo-parietal cluster: 200-220ms, peak at 208ms, *t*=5.33, *p*=0.003; see Table 2 and Figure 4).

**Table 2.** Significant clusters of activation for interaction effects (mismatch × stability) on ERPs in study 1. Columns are organized as in Table 1.

| **Study 1: mismatch × stability** | cluster | $x[mm]$ | $y[mm]$ | $z[ms]$ | $t_{66}$ | $Z_{\equiv}$ | $p_{FWE}$ | $k_E$ | $tw_{sig}$ [ms] |
| --- | --- | --- | --- | --- | --- | --- | --- | --- | --- |
| **A** stable MMN > volatile MMN | 1 | 17 | -19 | 204 | 5.73 | 5.14 | 0.001 | 591 | 180 - 220 |
|  |  | -17 | -25 | 196 | 5.68 | 5.10 | 0.001 |  |  |
| **B** volatile MMN > stable MMN | 1 | 42 | -78 | 200 | 5.63 | 5.07 | 0.001 | 119 | 188 - 236 |
|  |  | 55 | -68 | 200 | 5.17 | 4.72 | 0.005 |  |  |
|  |  | 64 | -62 | 200 | 5.04 | 4.62 | 0.007 |  |  |
|  | 2 | -60 | -57 | 208 | 5.33 | 4.85 | 0.003 | 29 | 200 - 220 |

**Figure 4.**
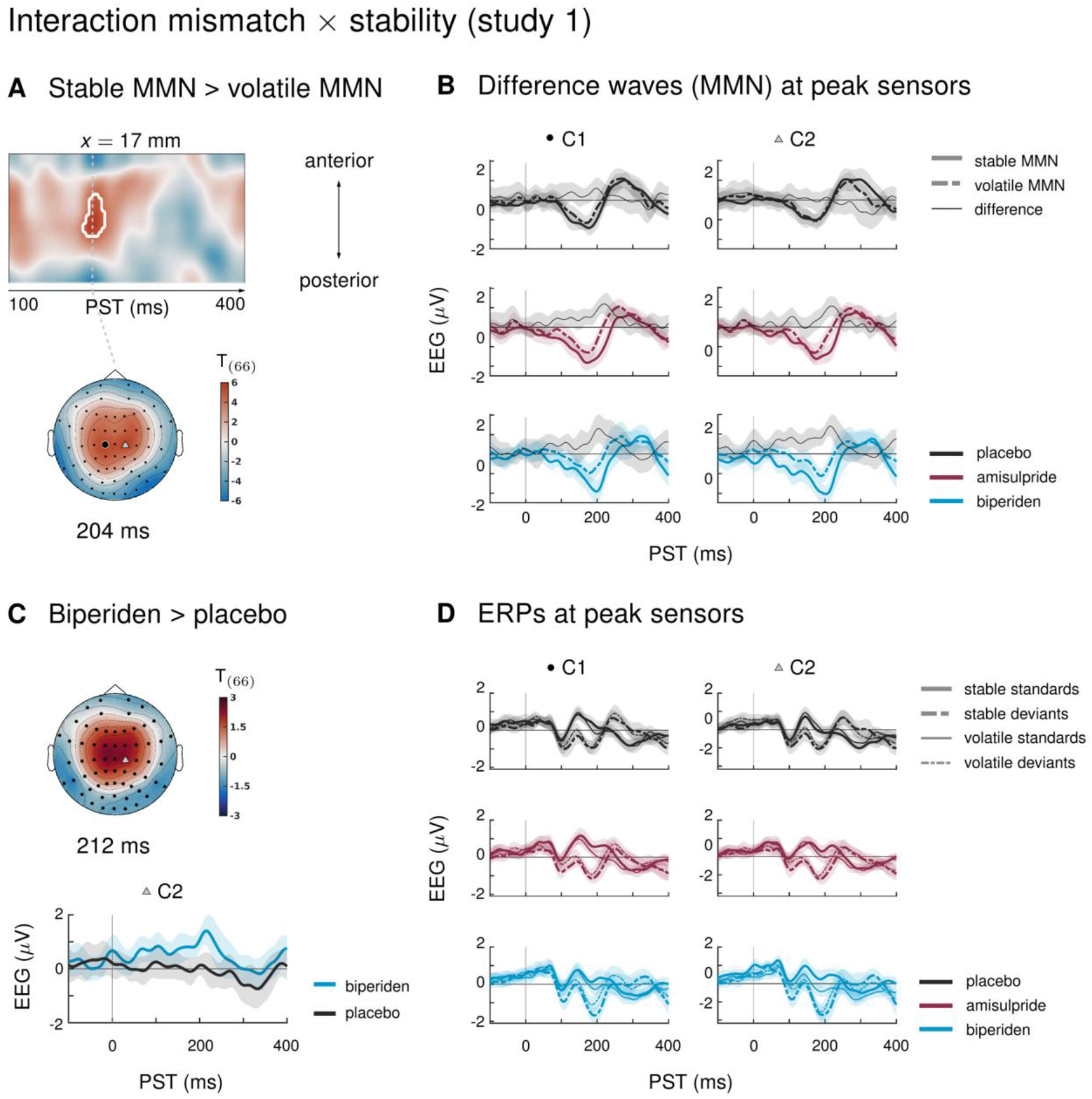
Interaction effects between stability and mismatch in study 1. **A** Regions of the time × sensor space where ERPs to tones in stable phases were more positive than ERPs to tones in volatile phases. Logic of display as in Figure 2. **B** ERP difference waves (standards − deviants) at the peak sensors for the two clusters shown in panel A, separately for the three drug groups. **C** Pharmacological effect on the interaction: at right central channels, the biperiden group showed a stronger interaction effect between mismatch and stability than the placebo group (significant only with application of spatio-temporal mask, see main text). Displayed is the *t*-map of the contrast and the difference waves (volatile – stable MMN) at sensor C2. **D** ERPs to standards and deviants at the same sensors as plotted in A and B.

Interaction effects in central channels reflected the following pattern: responses to standard tones were more positive and responses to deviant tones more negative during stable phases than during volatile phases (Figure 4D). The opposite was true for interaction effects at temporo-parietal clusters.

#### Interaction mismatch × stability × drug

In the ERPs at the sensors within the above clusters, it appeared that the interaction effect was mainly driven by the biperiden group (Figure 4B). Indeed, when examining the drug groups separately, the interaction effect was significant only in the biperiden group (208ms, *t*=4.96, *p*=0.009) at right central channels, but not in the placebo or the amisulpride group. However, there were no clusters for the three-way interaction with drug group which survived multiple comparison correction across the whole time × sensor space. The same held when zooming in on those clusters that showed significant interaction effects, using the functionally defined mask of the average interaction effects. However, focusing on only those parts of the time × sensor space where there was a significant positive interaction between mismatch and stability (cluster 1 in Table 2), we did find a significant three-way interaction such that the interaction of mismatch and stability was stronger in the biperiden group compared to the placebo group at 212ms at right central sensors (*t*=3.18, *p*=0.034 after small volume correction, see Figure 4C). Note that, similar to the constrained mask for the overall mismatch effects, this constrained mask was not part of our a priori analysis plan.

#### Drug effects on stable mismatch and volatile mismatch

In line with our analysis plan, we also examined the interaction of drug with mismatch during stable phases separately from mismatch during volatile phases. Because mismatch effects were stronger during stable periods of the experiment (see above: ‘Interaction mismatch × stability’), we suspected that we might also be more sensitive to the effects of the pharmacological manipulation in these periods.

Indeed, while there were no significant effects of drug group on mismatch in volatile phases, drug groups did differ significantly in their mismatch response during stable periods. Again, as for overall mismatch, pre-frontal sensors showed significantly reduced mismatch responses between 160ms and 168ms after tone onset in the biperiden group compared to the amisulpride group, peaking at 164ms (*t*=4.74, *p*=0.016). Additionally, later mismatch responses were significantly larger in the biperiden group compared to placebo at right central and centro-parietal sensors (see Table 3 and Figure 3B), again reflecting a delayed mismatch response under biperiden with a shift in topography from left frontal and pre-frontal towards right central and centro-parietal channels.

**Table 3.** Significant clusters of activation for pharmacological effects on stable mismatch in study 1. Columns are organized as in Table 1.

| **Study 1: Stable Mismatch** | cluster | $x[mm]$ | $y[mm]$ | $z[ms]$ | $t_{66}$ | $Z_{\equiv}$ | $p_{FWE}$ | $k_E$ | $tw_{sig}$ [ms] |
| --- | --- | --- | --- | --- | --- | --- | --- | --- | --- |
| **A PLA > BIP** | 1 | -26 | 56 | 204 | 4.56 | 4.24 | 0.027 | 8 | 204 - 212 |
|  | 2 | -26 | 61 | 224 | 4.36 | 4.07 | 0.049 | 1 | 224 - 224 |
| **B BIP > PLA** | 1 | 21 | -19 | 212 | 4.92 | 4.52 | 0.009 | 78 | 200 - 220 |
| **C AMI > BIP** | 1 | 17 | 72 | 164 | 4.74 | 4.38 | 0.016 | 11 | 160 - 168 |

When constraining the search volume using the average effect of stable mismatch, the delayed mismatch in the biperiden group was additionally significantly stronger than in the amisulpride group at 204ms in left pre-frontal sensors (*t*=4.17, p=0.041). Overall, the effects of biperiden on stable mismatch resembled the ones on overall mismatch signals, but with higher effect sizes, while there were no significant pharmacological effects on volatile mismatch.

## Results: Study 2

### Behavior in distraction task

Participants reacted to visual targets on average after 460.4ms (SD=54.2) and responded correctly to 95.6% (SD=5.3) of the presented targets. Reaction times did not differ significantly between drug groups (*F*=0.55, *p*=0.58), but there was a significant effect of drug group on hit rates (χ^2^=8.36, *p*=0.01, Figure 1D). Post-hoc pairwise tests indicated that hit rates in the galantamine group were significantly higher than in the levodopa group (*p*=0.019; the difference to the placebo group failed to reach significance: *p*=0.06). This result also held when excluding the participant with a hit rate below 75% (now placebo N=25; χ^2^=8.36, *p*=0.018; galantamine>levodopa *p*=0.017; galantamine>placebo *p*=0.104).

### ERP effects

#### Main effect of mismatch and interaction mismatch × drug

We found a strong effect of mismatch, where ERPs to standard tones were significantly more positive than ERPs to deviant tones from 100ms to 216ms after tone onset in a large cluster of frontal, fronto-central, and central sensors (peak at 176ms, *t*=14.13, p<0.001), and the opposite effect at left temporo-parietal and parietal sensors (100ms to 216ms, peak at 172ms, *t*=13.97, p<0.001). Standard and deviant ERPs were significantly different in nine additional clusters, which are listed in Table 4 and partly displayed in Figure 5.

**Table 4.** Significant clusters of activation for effects of mismatch (standards versus deviants) in study 2. Columns are organized as in Table 1.

| **Study 2: Mismatch** | cluster | $x[mm]$ | $y[mm]$ | $z[ms]$ | $t_{73}$ | $Z_{\equiv}$ | $p_{FWE}$ | $k_E$ | $tw_{sig}$ [ms] |
| --- | --- | --- | --- | --- | --- | --- | --- | --- | --- |
| **A standards > deviants** | 1 | 13 | -9 | 176 | 14.13 | Inf | 0.000 | 7583 | 100 - 216 |
|  |  | 4 | 18 | 160 | 13.76 | Inf | 0.000 |  |  |
|  |  | 42 | -25 | 124 | 10.85 | Inf | 0.000 |  |  |
|  | 2 | 0 | -9 | 396 | 8.66 | 7.16 | 0.000 | 1338 | 364 - 400 |
|  | 3 | 4 | -95 | 292 | 7.34 | 6.33 | 0.000 | 713 | 244 - 332 |
|  |  | 26 | -89 | 280 | 6.66 | 5.87 | 0.000 |  |  |
|  |  | 47 | -62 | 252 | 5.71 | 5.18 | 0.001 |  |  |
|  | 4 | 4 | 61 | 288 | 6.14 | 5.50 | 0.000 | 320 | 256 - 304 |
|  |  | -4 | 56 | 268 | 6.07 | 5.44 | 0.000 |  |  |
|  | 5 | -47 | -68 | 256 | 5.93 | 5.34 | 0.000 | 364 | 232 - 284 |
|  |  | -60 | -57 | 260 | 5.65 | 5.13 | 0.001 |  |  |
| **B deviants > standards** | 1 | -42 | -73 | 172 | 13.97 | Inf | 0.000 | 5637 | 100 - 216 |
|  |  | 55 | -68 | 196 | 10.87 | Inf | 0.000 |  |  |
|  |  | -42 | -73 | 124 | 10.38 | Inf | 0.000 |  |  |
|  | 2 | -8 | -30 | 256 | 7.41 | 6.38 | 0.000 | 2876 | 232 - 328 |
|  |  | -26 | -14 | 288 | 6.83 | 5.99 | 0.000 |  |  |
|  |  | 8 | -9 | 304 | 6.67 | 5.87 | 0.000 |  |  |
|  | 3 | 38 | -68 | 400 | 6.65 | 5.86 | 0.000 | 302 | 372 - 400 |
|  | 4 | 4 | 72 | 388 | 6.03 | 5.41 | 0.000 | 168 | 368 - 400 |
|  | 5 | 68 | 18 | 192 | 5.36 | 4.91 | 0.002 | 20 | 168 - 204 |
|  | 6 | -60 | -9 | 400 | 5.26 | 4.83 | 0.003 | 153 | 388 - 400 |
|  |  | -34 | -62 | 396 | 5.04 | 4.65 | 0.005 |  |  |

**Figure 5.**
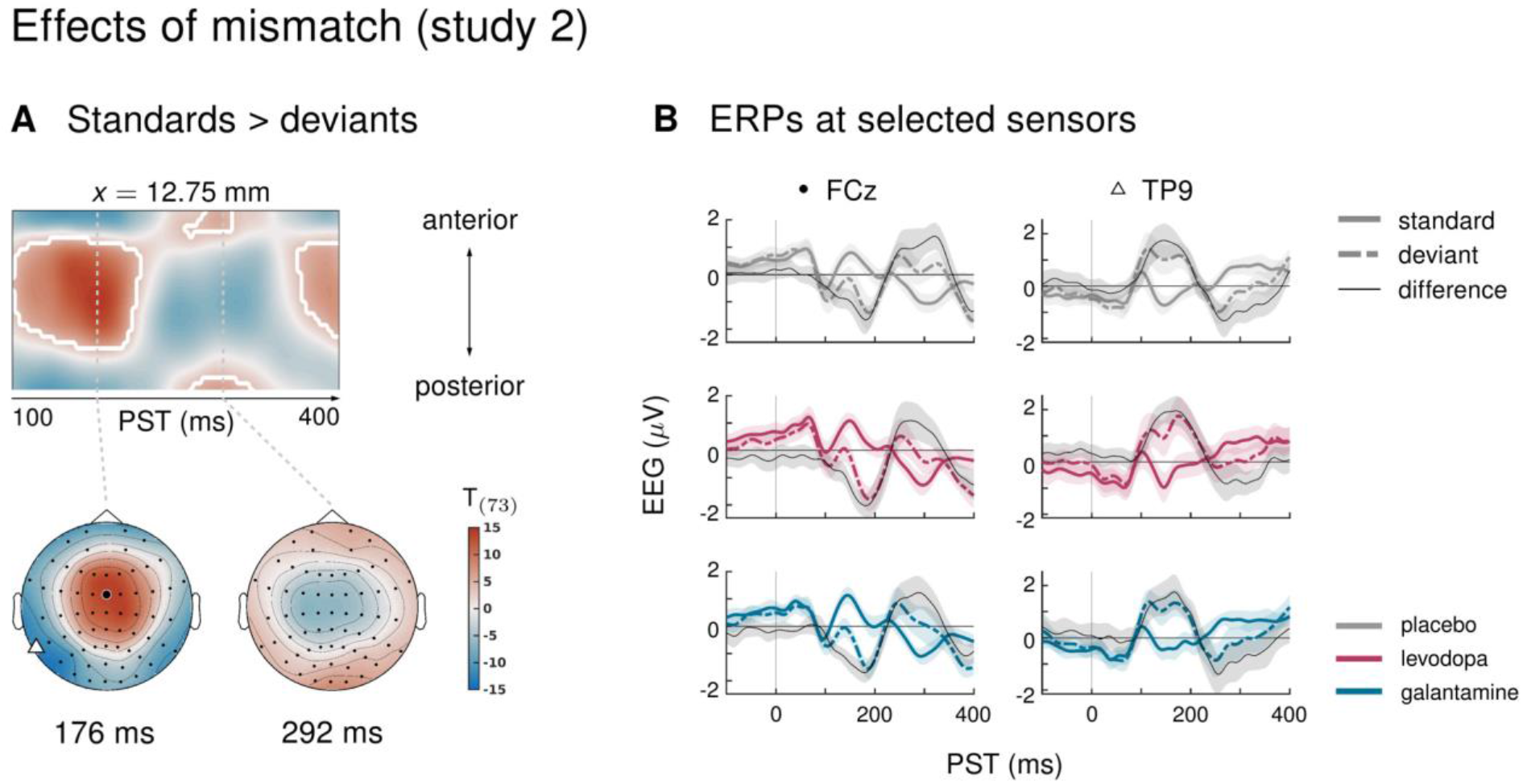
Main effect of mismatch in study 2. **A** Regions of the time × sensor space where ERPs to standards were more positive than ERPs to deviants. Logic of display as in Figure 2. As in study 1, mismatch was significant in large parts of the time × sensor space, including the classical mismatch negativity. **B** ERPs and difference waves for selected sensors, separately for the three drug groups. Mismatch signals were highly similar across all conditions (see main text).

There were no significant differences in mismatch ERPs between drug groups, both when considering the whole time × sensor space and when constraining the search volume to the significant average mismatch effect using the functionally defined mask. This also held when, by the same argument as in study 1, constraining the search even further by considering as a functional mask only the large cluster of frontal, fronto-central and central sensors described above, which corresponded to the classical time windows and sensor locations for the mismatch negativity.

#### Interactions mismatch × stability, and mismatch × stability × drug

Averaging over drug groups, ERPs showed no significant interaction between the factors mismatch and stability. In other words, mismatch responses did not differ between stable and volatile periods of the experiment. The three drug groups also did not differ in how mismatch ERPs were (not) affected by the stability of the current context (three-way interaction with factor drug). Examining mismatch responses in stable and volatile phases separately did not reveal any significant interactions with the factor drug either.

## Discussion

Above, we presented results from two pharmacological EEG studies which were designed to test the sensitivity of a new auditory oddball paradigm to cholinergic and dopaminergic modulations of synaptic plasticity. In study 1, we found that biperiden, a selective muscarinic (cholinergic) M1 receptor antagonist, delays and topographically shifts the mismatch negativity in this paradigm, while inhibiting dopaminergic transmission by administration of amisulpride did not affect mismatch-related ERPs. Neither elevated cholinergic nor dopaminergic transmission, as induced in study 2 by galantamine and levodopa, respectively, resulted in observable changes to deviance processing in our task.

Our paradigm allowed to examine processing of auditory deviants in two different contexts: during stable phases of the experiment, one tone reliably served as the ‘deviant’ (i.e., the unlikely) event, and the other as the ‘standard’. During volatile phases, the roles of standard and deviant switched more rapidly, requiring faster updating of the internal model of the acoustic environment. We found that antagonizing muscarinic cholinergic receptors with biperiden affected deviance processing particularly during stable phases of our task and led to a significant interaction between deviance and stability.

### Delayed and topography-shifted mismatch responses under biperiden

In study 1, mismatch responses in the biperiden group peaked later and were distributed more towards right centro-parietal channels than in the other drug groups (Figure 3D). This resulted in significantly smaller mismatch amplitudes (reduced MMN) at pre-frontal sensors early on, in classical MMN time windows (biperiden vs. amisulpride), and significantly larger MMN at centro-parietal sensors later (biperiden vs. placebo).

Effects of cholinergic agents on MMN have been demonstrated repeatedly, mostly showing enhanced mismatch amplitudes or shortened MMN latencies in response to stimulation of nicotinic cholinergic receptors (Baldeweg et al., 2004; Harkrider and Hedrick, 2005; Dunbar et al., 2007; Martin et al., 2009; Knott et al., 2012; Hamilton et al., 2018). In contrast, previous investigations of the effects of antagonizing muscarinic cholinergic receptors have yielded less consistent results. Studies using the muscarinic antagonist scopolamine have reported reductions of MMN amplitudes (Pekkonen et al., 2001), no effects on MMN (Pekkonen et al., 2005), and reduced P300 responses to targets in active oddball tasks (Meador et al., 1989; Curran et al., 1998; Brown et al., 2015). Here, we used a passive auditory oddball task, following the recommendation that MMN assessment is optimal when the participant’s attention is directed away from the auditory domain (Näätänen, 2000), and tested the effects of biperiden.

Biperiden differs from scopolamine in the specificity of its binding affinity: it has about tenfold higher affinity for M1 as compared to M2–M5 receptors (Bolden et al., 1992). Two studies have tested the effects of biperiden on deviance detection in passive auditory oddball tasks (Klinkenberg et al., 2013; Caldenhove et al., 2017). Neither study found effects on MMN, but hints of a potential effect on P3a amplitudes. Importantly, both used only half the dose (2mg) of biperiden as administered here, which might explain the difference in findings compared to our study.

Most previous pharmacological studies of MMN restricted their examination of drug effects on MMN to specific sensors and time points, mostly based on average MMN difference waves. Here, we provide a characterization of the drug effect across the full time × sensor space, while correcting for multiple comparisons across time and sensors. This analysis revealed both a delay in peak MMN amplitude, and a shift in topography in the biperiden group (Figure 3). Importantly, this shift affects traditional MMN sensors (Fz, FCz, Cz), which have mostly been examined in previous studies, less than those at the border of the MMN scalp distribution (Fp1, Fp2, C2, C4), which is where we found significant effects of biperiden. Another strength of our study design – with total N=162 across studies – was the use of individual drug plasma level estimates in the group level GLM, based on the analysis of blood samples, which allowed us to account for interindividual differences in pharmacokinetics. We further controlled for potential confounds by means of our inclusion criteria, e.g., excluding smokers to avoid effects of baseline nicotine levels. The focus on male participants was intended to avoid confounds of fluctuating estrogen levels, which have been found to significantly impact on dopaminergic and cholinergic systems (Gasbarri et al., 2012; Colzato and Hommel, 2014; Barth et al., 2015). However, this also constitutes a significant limitation of our study, as it means that our results may not equally apply to both sexes and will therefore need to be replicated in a more representative sample in future work.

Surprisingly, in study 2, we did not find an effect of galantamine on mismatch responses. This is in contrast to a previous report showing an augmentation of MMN under the same dose of galantamine as administered here (Moran et al., 2013). The study by Moran and colleagues employed a ‘roving’ oddball paradigm (Garrido et al., 2008): a tone sequence comprised mini-blocks of 6-10 tone repetitions, where consecutive mini-blocks differed in frequency, and the first tone of a block represented the deviant. Importantly, in their paradigm, every deviant indicated the onset of a new context, and contexts (mini-blocks) lasted for at least six tones. In contrast, in our paradigm, mini-blocks of repeated tones tended to be much shorter (less than 20% of mini-blocks with 6 repetitions or more, more than 60% consisting of 2 repetitions or less), making our paradigm considerably more volatile overall. This comparably high tonic volatility could mean that precision-weights on PEs were already high; this might have prevented any increase in sensory precision afforded by galantamine to be expressed in the mismatch ERPs in our study due to a ceiling effect.

Notably, our analysis differed from previous investigations of the effects of biperiden and galantamine on the MMN (e.g., (Klinkenberg et al., 2013; Moran et al., 2013; Caldenhove et al., 2017)) in terms of pre-processing choices such as the amount of high-pass filtering and the application of baseline correction. To examine the robustness of our main results to these analysis choices and to rule out that differences in results between studies were simply due to different pre-processing strategies, we re-analyzed our data using equivalent settings to these previous reports. The results of this analysis, presented as part of the supplementary material (section 2.4), were largely supportive of our claims: we also found significant effects of biperiden (versus placebo and amisulpride) on mismatch signals that were compatible with an increase in MMN latency, even when adopting the same pipeline as used in previous studies (see section 1.4 of the supplementary material for details on this pipeline and section 2.4 for the results). Similarly, there was still no effect of galantamine on mismatch responses in a whole time × sensor space analysis based on this adapted pipeline.

### Biperiden and the influence of environmental volatility on mismatch processing

In classical oddball paradigms, the occasional deviant represents a rule violation, but its impact on subsequent rule representation is limited, as the tone sequence reverts back to the standard tone immediately. In contrast, the roving oddball paradigm examines model *updating* in a changing environment, as every deviant signals the onset of a new rule. In our paradigm, the relevance of the detected rule violation to the representation of the rule additionally varies across different periods of the experiment: during more stable phases, oddballs represent noise and deviants should not lead to a major update of the current belief about the underlying rule. In contrast, during volatile phases, the probabilities of the two tones sometimes reverse and deviants thus occasionally signal the onset of a new rule. Theoretical treatments suggest that this volatility can impact on the size of belief updates in two opposing ways (Mathys et al., 2011). On the one hand, increased belief uncertainty due to environmental volatility should increase learning rates (i.e., belief updates) – in other words, deviants are more meaningful in volatile phases due to the occasional rule switch. On the other hand, stable phases allow for a more precise prediction of the input than volatile phases, as beliefs about the more likely tone occurrence are allowed to accumulate for longer. This suggests an increased impact of deviants during stability. It is a priori not clear which of these two opposing effects would dominate in a given setting. In our case, we examined this question by contrasting mismatch effects between stable and volatile periods of our task.

In study 1, we found a significant difference between stable and volatile mismatch responses, such that mismatch was stronger in stable than in volatile periods. This mirrored previous reports of volatility effects on mismatch signals (Todd et al., 2014; Dzafic et al., 2020). However, in our study, this was mainly due to the altered mismatch response in the biperiden group, which was particularly affected during stable mismatch. Neither the placebo group nor the amisulpride group showed interaction effects on their own, and, when directly contrasting the groups and focusing on the cluster of central channels that showed the average effect across drug groups, the effect was significantly stronger in the biperiden group compared to placebo. No significant differences between stable and volatile mismatch responses were found in study 2. Note however, that under a different pre-processing pipeline (with more aggressive correction of slow drifts) and/or trial definition (optimized for retaining more trials per condition), we do see these interaction effects across all drug groups in both studies (supplementary material section 2.4).

It should also be noted that previous reports have presented effects of block-wise volatility changes on mismatch processing in single-channel analyses (focusing only on Fz, (Todd et al., 2014; Dzafic et al., 2020), and that the whole-volume corrected effect presented in (Dzafic et al., 2020) did not replicate in a validation data set, suggesting that the effects of block-wise volatility on mismatch might be relatively subtle compared to the size of the mismatch effect itself (and thus require more aggressive pre-processing or higher trial numbers to be robustly detected). This subtlety of block-wise stability variation might also be explained by the two opposing effects of volatility on precision-weights described above, which might cancel each other to some degree.

Instead of averaging EEG responses within different stimulus categories (‘standard’, ‘deviant’) or within different blocks (‘stable’, ‘volatile’), we have previously capitalized on the history-dependence of EEG amplitudes in learning paradigms such as the MMN, where trial-wise amplitude changes carry information about the temporal dynamics of the belief updating process (Lieder et al., 2013a; Stefanics et al., 2018; Weber et al., 2020). This is particularly attractive when considering the impact of environmental volatility on learning rates (Behrens et al., 2007; Mathys et al., 2011). However, in the absence of a forward model whose inversion allows for inferring beliefs directly from EEG data, these analyses are restricted to ideal observer analyses (Stefanics et al., 2018; Weber et al., 2020) due to the passive nature of the paradigm. In the supplementary material (sections 1.4 and 2.4) we present such an analysis for the current data set, the results of which were highly consistent with our previous reports applying the same model to other MMN paradigms (Stefanics et al., 2018; Weber et al., 2020), in that we found multiple, hierarchically related prediction errors underlying EEG mismatch signals: trial-wise EEG amplitudes of the classical MMN component correlated with lower-level prediction errors about tone probabilities, while later P3-like components scaled with a higher-level prediction error about environmental volatility.

Importantly, in line with the results from the conventional averaging approach, we find that only biperiden affected the EEG signatures of model-derived precision-weighted prediction errors – neither dopaminergic manipulations nor galantamine showed significant differences to the placebo group. Moreover, the biperiden effect concerned not only signatures of low-level prediction errors, reflecting mismatches between expected and actual tone identities, but also signatures of higher-level prediction errors serving to refine beliefs about environmental volatility. However, in the main text of the paper, for ease of accessibility and to enable direct comparison with previous work on dopaminergic and cholinergic effects on the MMN, we have focused our conclusions on the results of the conventional ERP analysis.

### Future directions

In this study, we employed a conventional ERP analysis, but considered all sensors and time points under multiple comparison correction, to detect effects of experimental conditions that manifest as differences in evoked response amplitudes within our time-window of interest.

Our pattern of results – an apparent biperiden-induced shift in mismatch responses from an early to a later peak, and from frontal to central channels – suggests that methods which exploit the rich temporal information in the EEG signal more than the amplitude-based approach could help us to further understand the impact of cholinergic neurotransmission on perceptual inference in our task. Examples for this are principal component analysis (PCA) based analyses (Hunt et al., 2015), which take into account the topography as well as the time course of the ERP, or dynamic causal modeling (DCM), which interprets scalp-level effects in terms of extrinsic (between-area) connectivity changes and local effects (such as synaptic gain modulation within an area) in an underlying network of sources (David et al., 2006; Kiebel et al., 2006; Garrido et al., 2007). Future analyses of the current data set might employ this technique to infer on low-level (synaptic) mechanisms underlying the observed pharmacological effects, e.g., biperiden-induced changes in post-synaptic gain of supragranular pyramidal cells in auditory cortex (Moran et al., 2013; Schöbi et al., 2021).

The current analysis demonstrates the sensitivity of our paradigm to muscarinic receptor status. In contrast, and in line with previous reports (Kähkönen et al., 2002; Leung et al., 2007, 2010; Korostenskaja et al., 2008), dopaminergic challenges in both of our studies did not affect mismatch responses. This differential sensitivity to cholinergic versus dopaminergic neuromodulation may prove valuable for understanding and predicting differential treatment responses in individuals diagnosed with schizophrenia. Importantly, while the reduction of MMN amplitudes in patients compared to healthy controls is robust and of large effect size (Erickson et al., 2016), there is still considerable inter-individual variation in MMN amplitudes among patients (Light and Swerdlow, 2015), supporting the idea that different subgroups of patients might differ in their MMN expression. Based on our results, reduced MMN in patients might be relatively more indicative of cholinergic versus dopaminergic dysregulation of synaptic plasticity. Critically, subgroups with differences in muscarinic receptor availability have been reported (Scarr et al., 2009), consistent with the possibility that the differential contribution of acetylcholine versus dopamine to NMDAR dysregulation represents a key pathophysiological dimension to explain clinical heterogeneity among patients with schizophrenia (Stephan et al., 2006, 2009).

Distinguishing these potential subtypes of schizophrenia could be highly relevant for treatment selection, as some of the most effective neuroleptic drugs (e.g., clozapine, olanzapine) differ from other atypical antipsychotics (e.g., amisulpride) in their binding affinity to muscarinic cholinergic receptors. To establish the utility of our paradigm in the clinical context, prospective patient studies are needed, which test whether this readout of cholinergic neurotransmission is predictive of treatment success in individual patients. In particular, such a prediction may become possible by adopting the “generative embedding” strategy frequently used in translational neuromodeling and computational psychiatry (Stephan et al., 2017): this involves estimating synaptic variables of (generative) neuronal circuit models of MMN and using these estimates as features for subsequent machine learning. While the potential of this computational strategy, in the specific context of muscarinic manipulations of the MMN, was demonstrated by a recent rodent study (Schöbi et al., 2021), an important question for future work is whether it can be successfully translated to a clinical setting.

## Acknowledgements

This study was supported by the University of Zurich (KES) and the René and Susanne Braginsky Foundation (KES). We thank Diana Kutyniok (MPI for Metabolism Research Cologne) for performing DNA isolation and SNP-genotyping.

