## Supplementary Material for "Auditory mismatch responses are differentially sensitive to changes in muscarinic acetylcholine versus dopamine receptor function"

#### Overview

1. Supplementary Methods
    1. Specification of the drug plasma level covariate
    2. Blood analysis genetic variation
    3. GLM specification for genetic effects
    4. **Robustness of our main results to analysis choices**
2. Supplementary Results
    1. Main effect of stability and interaction stability with drug group
    2. Control analyses: Excluding datasets based on the distraction task
    3. Genetic effects and pharmaco-genetic interactions
    4. **Robustness of our main results to analysis choices**

### 1 Supplementary Methods

#### 1.1 Specification of the drug plasma level covariate

In our group-level GLMs, we introduced a covariate for the estimated drug plasma concentration levels of both pharmacological agents, where we allowed for an interaction with the drug factor and mean-centered the covariate within drug groups. This section summarizes our motivation for this covariate specification in a design with multiple drug groups, where the main effect of interest is the (mean) difference between groups (or drugs) on brain activity.

A covariate coding for drug plasma levels can have two purposes. On the one hand, it can be used to explain (noise) variance, i.e., physiological or cognitive effects of the drug that depend on its dose and are not related to our task effect of interest but do change the measured brain signal. This means using it as a covariate of no interest (or a confound). On the other hand, one might use the readout as a proxy for the amount of drug that reached the brain and exerted its influence on the processes of interest there (although the relationship between peripheral drug plasma levels and drug concentration in the brain is non-trivial). In that case, it is a covariate of interest, and one might want to explicitly test for dose-dependent effects in our EEG signals.

##### Mean-centering the covariate

Guidelines in educational resources for neuroimaging analyses usually recommend to not mean-center continuous covariates within different groups. For example, in an educational blog entry http://mumford.fmripower.org/mean_centering/, Jeanette Mumford notes the following:

«Also important is that if you have multiple groups (...) you should not mean center within each group separately. The reason for this is that your continuous covariate may actually describe some of the group differences, and then mean centering within group will remove this important aspect of the covariate. For example, if you have two groups and one is significantly younger than the other, this difference in ages between groups may explain brain activation differences. If you mean center age within each group, then each group's set of ages will be centered about the same value, 0, and then you risk detecting group differences that are actually just attributable to age.»

In this example, age is a confound for group differences. In such a case, if the covariate codes for a confound unrelated to the group factor, and there are mean differences between groups in this variable, any difference in brain activity between groups could equally be explained by the group factor and the covariate. In these situations, it is especially important to allow the covariate to explain this difference to avoid false positives for the group factor.

This is fully consistent with other recommendations, e.g., the Wiki pages for the fMRI analysis software FSL (https://fsl.fmrib.ox.ac.uk/fsl/fslwiki/GLM#Two-Group_Difference_Adjusted_for_Covariate).

However, differences in mean drug plasma level between a placebo condition and any pharmacological manipulation are not (merely) confounds, but instead constitute the group difference we want to test for (using the factor group).[1] (If one were interested in the effects of age on brain activity, one would hardly form two age groups, only to regress out the effect of ‹age› as an un-centered covariate.) In our pharmacological design, the confound variables are the peripheral (e.g., blurred vision), and indeed central (e.g., drowsiness, vertigo, etc.) side-effects of our drugs. These covary with the peripheral drug plasma level. However, the effects of interest on the central nervous system (e.g., altered auditory mismatch processing) may also be a function of the plasma level. Consequently, the fact that dose-dependent effects (and subsequently group differences) in our measurement readout (mismatch-related ERPs) can be explained both by our intended experimental manipulation (e.g., levels of acetylcholine in cortical areas responsible for auditory processing), and by potential side-effects of the drugs, both peripherally and centrally, is a challenge inherent to our experimental design, where we administer the drugs peripherally.

This cannot be resolved simply by using drug plasma levels as a confound variable. Instead, in the future, (non-invasive) tools that give us direct control over the release of acetylcholine (ACh) into our areas of interest without influencing ACh levels elsewhere in the nervous system, will provide definite answers to these questions. Meanwhile, in our current analyses, we use the covariate ‹drug plasma level› only to only model the slope (dose-dependent effects) within each drug group, but never the mean difference between groups – this is what the factor group is for.[2]

---

[1] Strictly speaking, this only holds for comparing the placebo group to a drug group. However, for comparing two drugs with each other, it still makes sense to mean-center within groups, as the difference in dose between the drugs is most likely not meaningful (the same dose means different things for different drugs).

[2] Note that mean-centering the covariate ‹drug plasma level› in each group separately, still allows testing for dose-dependent effects explicitly by using an *F* - or *t*-contrast on the covariate (the result will be the same as in the un-centered case). However, the concentration of drug in our target brain area may differ considerably from the concentration in the peripheral blood. Because of this uncertainty, in the current analysis presented in the main text, we refrained from investigating dose-dependent effects using the drug plasma level covariate and use it only to explain within-group variance.

#### Interaction with factor drug group

We entered the drug plasma level values in separate covariates, one per drug group (none for placebo), and thus estimated separate beta weights for them. In each covariate, the values within the other drug groups (and placebo) are all zero. This can be interpreted as allowing for an interaction of the covariate ‹drug plasma level› with the group factor. We do this, because having them as one covariate would ignore that the measurements for different drugs naturally live on different scales, and that the dose-effect relationship is expected to be very different across different drugs. It is thus questionable whether estimating a single beta for the linear effect of drug would be meaningful at all.

When testing for dose-dependent effects directly, it is important to note that interpreting a difference in group means is complicated in the presence of a significant interaction. For example, a section from the FSL Wiki reads (https://fsl.fmrib.ox.ac.uk/fsl/fslwiki/GLM#Two-Group_Difference_Adjusted_for_Covariate):

«Two Groups with continuous covariate interaction

(...) A test of the difference in group means does not make sense in the presence of a significant interaction, as the interaction indicates that the group difference varies as a function of age. Therefore, focusing on a single age is not likely all that interesting. Instead, if the interaction is not significant, the group mean differences can be obtained from the previous model. »

Again, our case (drug plasma levels) is different from their example (age), as it is directly related to the group factor: e.g., greater differences between drug groups at higher doses might be expected. Moreover, as stated above, we expect that the dose-effect relationship differs between drugs. Therefore, we never estimate a model without interaction. Our main question of interest is: at the doses at which participants received the drugs (which for all drugs, except placebo, resulted in some mean drug plasma level across participants which is larger than zero), do these drugs have dissociable influences on brain activity? The mean drug plasma level is not an arbitrary point of comparison here (in contrast to the mean age of a randomly assigned experimental group) but depends on our careful choice of the dose at which our drugs were administered. Acknowledging that most drugs have complex, non-linear dose dependent effects on brain processes, the experimenter chooses a dose for which previous research has provided sufficient evidence for the effect of interest being present while potential side-effects are minimized. Moreover, in the GLM, we always model these dependencies as linear, which can be seen as a local approximation to the otherwise non-linear true relationship. Within this local approximation (i.e., in proximity to the mean drug plasma level observed in our sample), a significant interaction of the covariate with the factor group merely captures the (expected) differential dose-effect relationships for different drugs.

#### 1.2 Blood analysis for genetic variation

We collected an additional blood sample per participant to assess genetic variation at functional single nucleotide polymorphisms (SNPs) of two genes relevant to the pharmacological intervention. For genetic analyses, 20 ml of blood were collected in tubes containing ethylene-diaminetetraacetic as anticoagulant, centrifuged at 10°C for 10min at 3000xg to separate the buffy coat and finally stored at -86°C until analysis. In this study, we assessed genetic variations at SNPs of the genes that code for the Catechol-O-methyl-transferase (COMT, using the rs4680 SNP) and the choline acetyltransferase (ChAT, using the rs1880676 SNP) enzymes.

Specifically, DNA from the buffy coat was isolated using the QIAamp DNA Blood Mini Kit (Cat No. 51106, Qiagen GmbH, Hilden, Germany) according to manufacturer's protocol. Concentration and quality of the DNA was assessed with a UV/Vis-spectrophotometer (ND-1000, Peqlab GmbH, Erlangen, Germany). Then, 20ng of DNA was analyzed in triplicates using allelic discrimination assays (TaqMan SNP Genotyping Assays, Applied Biosystems by Thermo Fisher Scientific Inc., Waltham, MA, USA). Genotyping PCR was performed on a 7900HT Fast Real-Time PCR system (Applied Biosystems) and the data analyzed with Sequence Detection Software (SDS) 2.3 (Applied Biosystems). DNA isolation and SNP-genotyping was performed by the Max Planck Institute for Metabolism Research in Cologne. These procedures were equivalent for both studies.

#### 1.3 GLM specification for genetic effects

In two additional GLMs per effect of interest, we considered variations in the expression of our experimental factors in EEG activity that are due to different variants (SNPs) of two genes that determine the availability of dopamine (DA) and ACh in the brain, respectively: COMT – which encodes catechol-O-methyltransferase, a key enzyme for the degradation of DA – and CHAT – which encodes choline acetyltransferase (abbreviated as ChAT), the enzyme responsible for synthesis of ACh. In one GLM, an additional covariate 'COMT polymorphism' coded for SNPs of the COMT gene, in the other GLM, an equivalent covariate 'CHAT polymorphism' was used. These covariates served to account for the possible contribution of genetic effects to individual variability in DA/ACh availability. They were used to test for direct effects on mismatch-related

EEG activations and for pharmaco-genetic interaction effects on brain activity during mismatch processing. In specifying these covariates, we allowed for an interaction with the drug factor, and mean-centered the covariates within drug groups.

We examined whether these covariates explained additional variance using *t*-tests for positive and negative effects of the covariates (for COMT: within placebo and amisulpride groups, for CHAT: within placebo and biperiden groups) and tested for pharmaco-genetic interaction effects using differential contrasts (COMT: placebo vs. amisulpride; CHAT: placebo vs. biperiden) in both directions. Again, we performed this analysis once across the whole time-sensor space, and subsequently investigated genetic effects within a smaller, functionally constrained search volume of significant average effects (orthogonal contrast). The functional masks were identical to the ones described for the main group-level GLMs.

In study 2, genotyping was inconclusive for two individuals for both gene variants (N=1: levodopa, N=1: galantamine), and additionally for one individual only for the COMT gene (N=1: levodopa). Therefore, the GLMs for CHAT effects in the EEG are based on a sample of N=76, and those for COMT effects are based on a sample of N=75.

#### 1.4 Robustness of our results to analysis choices

Following reviewer comments in a previous submission of our manuscript, we examined the robustness of our main findings to changes in analysis strategy. The concerns we aimed to address were mainly three:

1. Our **pre-processing choices** (average reference, very mild high-pass filtering, no baseline correction) in combination with our **whole time-sensor space analysis** (correcting for multiple comparisons across sensors and time points) might be too conservative, potentially leading to
    a. us being insensitive to pharmacological effects on mismatch responses in our data and
    b. some of the volatility effects (i.e., the interaction mismatch*stability) being obscured or even driven by slow drift artefacts.
2. Our **trial definition** results in a relatively low number of trials per condition. While we have chosen this definition due to its specificity, we also noted alternative trial definitions in our analysis plan which allow for a higher number of trials being retained for analysis. Again, the concern was that our choice might lead to a lack of power for

    a. detecting pharmacological effects on the MMN in our paradigm, and
    b. detecting differences between stable and volatile phases (as trial numbers must be split up again among these phases)
3. It was noted that we rely on a classical model-free ERP analysis even though we have previously analyzed and interpreted the MMN in terms of prediction errors in a hierarchical Bayesian inference scheme. The **model-based analysis** for the current data set was also outlined in the original analysis plan and was perceived to be a more appropriate way of examining effects of (trial-wise) volatility beliefs on mismatch responses.

To address these concerns for future readers of our manuscript, we performed the following additional analyses:

1. We examined the robustness of the *pharmacological effects* on mismatch under a **different pre-processing pipeline and a different trial definition**, using both our whole time-sensor space analysis approach, and a more sensitive **region-of-interest (ROI) approach** based on previous literature.
2. We also examined the robustness of the *interaction effect mismatch*stability* to the choice of pre-processing pipeline and trial definition.
3. We performed the **model-based single-trial analysis** outlined in our analysis plan and examined the effects of *drug group on the EEG correlates of hierarchical prediction errors*.

#### Alternative pre-processing and trial definition

To make our results more directly comparable with previous reports on the effects of biperiden and galantamine on the MMN (Klinkenberg et al., 2013; Moran et al., 2013; Caldenhove et al., 2017), and address concerns of being not sensitive enough, we *copied the pre-processing settings from these reports*, resulting in the following changes to our previous pipeline:

- a re-referencing to a linked mastoid reference
- a strong high-pass filter of 1 Hz
- a baseline correction using a 100ms pre-stimulus time window

All other pre-processing steps remained as reported in the main text.

Additionally, we used a different standard and deviant definition (that we had already specified as part of our analysis plan): instead of only considering tones as standards and deviants that followed at least 5 repetitions of the same tone (“roving” MMN definition), we instead considered every tone after at least 2 repetitions. Because this results in many more standard tones than deviant tones, we only used a subset of these standard tones, choosing them such that for a given number of preceding tone repetitions, there were as many standards as there were deviants (“fair” MMN definition, see also the description in the publicly available analysis plan).

Alongside our whole time-sensor space analysis, we also performed a region-of-interest (ROI) analysis, again *exactly copying the procedure used in previous reports* (Klinkenberg et al., 2013; Caldenhove et al., 2017). In particular, we focused on the three electrodes Fz, FCz, and Cz. We extracted for every participant the peak MMN amplitude and its latency. We defined the MMN peak as the minimum of the difference wave (ERP to standard tones minus ERP to deviant tones), obtained using the adjusted pre-processing pipeline and the “fair” trial definition, in the time window between 150ms and 250ms. MMN peak amplitudes and latencies per participant and electrode were entered into separate 3x3 ANOVAs (factor 1: drug group, factor 2: electrode) for each study separately. Significant main effects were followed up by post-hoc *t*-tests.

#### A model-based perspective on trial-by-trial auditory ERPs

We have previously modelled the trial-wise auditory ERPs in MMN paradigms using a hierarchical Bayesian model of belief updating (Mathys et al., 2011) and demonstrated that later mismatch responses reflect higher-level prediction errors involved in estimating the stability of the auditory environment (Weber et al., 2020). However, volatility effects on MMN *have also previously been demonstrated using a “contextual” approach* (contrasting phases with different stability levels) (Todd et al., 2014; Dzafic et al., 2020). In the main text, we followed this approach, and experimentally demonstrate an interaction between mismatch and stability, *supporting the pure model-based analyses* (Weber et al., 2020) that have operated under *constant volatility paradigms.*

Here, we support the findings presented in the main text by demonstrating the robustness of the blocked stability effects to different analysis pipelines (in particular, baseline correction, high-pass filtering, and trial definition). Moreover, we complement the block-wise model-free analysis by additionally reporting the results of a model-based analysis, which we had fully specified already in our analysis plan. For convenience, we include the paragraphs describing this analysis here again.

*Trial Definition*

For the single-trial analysis, we only defined one trial type (‘tone’), i.e., we kept all tone events, and we did not perform an average over trials.

*Conversion to Images and Smoothing*

We converted the final pre-processed file of each participant into 4D (scalp x within-trial time points x single trials) images using the SPM function spm_eeg_convert2images. Subsequently, we applied a spatial smoothing with a Gaussian kernel of 16 mm FWHM in both spatial (x and y) dimensions. These smoothed images entered a first level regression GLM per participant.

*Model*

In the absence of any behavioral responses of participants to the auditory input, we modeled all participants as surprise-minimizing ('Bayes-optimal') agents under the perceptual model of the 3-level hierarchical Gaussian Filter (HGF) for binary inputs. This means that we chose the perceptual parameters of this model such that an exposure to the tone sequence used in our task leads to least surprise over all inputs. To this end, we used the Matlab function tapas_fitModel from the HGF toolbox (v5.1.0), distributed as part of TAPAS (release v3.0.0) with the tapas_bayes_optimal_binary function as a pseudo response model. We previously examined the robustness of this parameter estimation for the given tone sequence with respect to changes in the prior means and uncertainties for each parameter. The results of this analysis are documented in DPRST_HGF_Bayes_optimal_parameters.pdf (available as part of the publicly available analysis plan). As a result, we used the belief trajectories of the agent characterized by the parameter values (and starting values) summarized in the last row of Table S1.

| ***Parameter*** | $\mu_2^{(0)}$ | $\mu_3^{(0)}$ | $\sigma_2^{(0)}$ | $\sigma_3^{(0)}$ | $\kappa_1$ | $\kappa_2$ | $\omega_2$ | $\omega_3$ |
|---|---|---|---|---|---|---|---|---|
| *Prior mean* | 0 | 1 | 0.25 | 1 | 1 | 1 | -3 | -10 |
| *Prior variance* | 0 | 0 | 0 | 0 | 0 | 0 | 16 | 0 |
| *Posterior mean* | 0 | 1 | 0.25 | 1 | 1 | 1 | -3.03 | -10 |

Table S1: Parameter settings for the HGF. The first two rows show the priors under which the surprise-minimizing parameters were estimated, the last row displays the result of the estimation and thus the values used in the simulation of belief trajectories. All perceptual parameters and starting values of beliefs were fixed to their prior means (indicated by zero prior variance) except for the tonic learning rate on the second level, ω2.

From the belief trajectories of this agent, we extracted the trial-wise estimates of precision-weighted prediction error on two levels of the hierarchy, $abs(\varepsilon_2)$ and $\varepsilon_3$, corresponding to PEs about stimulus occurrences and learning signals for the estimation of environmental volatility, respectively. These vectors entered the first level GLM of trial-wise EEG signals for each participant as multiple regressors.

*First Level GLMs*

Our model-based vectors of precision-weighted PEs served as regressors in a general linear model (GLM) of trial-wise EEG signals for each participant separately, correcting for multiple comparisons over the entire time-sensor matrix, using Gaussian random field theory. We did not orthogonalise the regressors. We used the same regressors for all participants but excluded entries for trials that were rejected during EEG preprocessing to ensure the regressors have the same length as the number of available EEG data trials for each participant. On the first level, we used a multiple regression model and defined separate *t*-tests for each regressor to examine positive and negative correlations of EEG amplitudes with each regressor. This GLM only serves to provide the beta images used for the second level analysis, therefore, we will not threshold the images based on the significance of the tests under either peak- or cluster-level familywise error (FWE) correction.

*Group Level GLMs*

The GLMs and tests on the second level for the model-based analysis were specified exactly as the GLMs for the group level conventional ERP analysis: separate GLMs for each computational quantity, which implement a factorial design with the between-subject factor 'Drug group', a covariate for drug plasma levels, and the same second level *t* contrasts for each computational quantity as in the conventional ERP analysis for each contrast of interest (see main text).

*Visualization as ERPs*

To visualize the effects of our model-based analysis in terms of ERPs, we used an additional trial definition which averaged the ERPs for the 10% lowest prediction error trials ("standards") and the 10% highest prediction error trials ("deviants") and computed the grand averages across participants and difference waves for these ERPs.

### 2 Supplementary Results

#### 2.1 Main effect of stability and interaction stability × drug group

##### Study 1: Main effect of stability

ERPs in three clusters showed a significant main effect of stability (when averaging over standard and deviant tones and the three drug groups): early on, pre-frontal (144-148ms, peak at 148ms,

$t$=4.76, $p$=0.017) and occipital sensors (136-156ms, peak at 144ms, $t$=5.04, $p$=0.007) showed stability effects; later on, stability affected ERPs at parietal sensors (272-292ms, peak at 284ms, $t$=5.32, $p$=0.003; see Table S1 and Figure S1). In all three clusters, ERPs to tones in stable phases were significantly more positive than ERPs to tones in volatile phases (see Figure S1B).

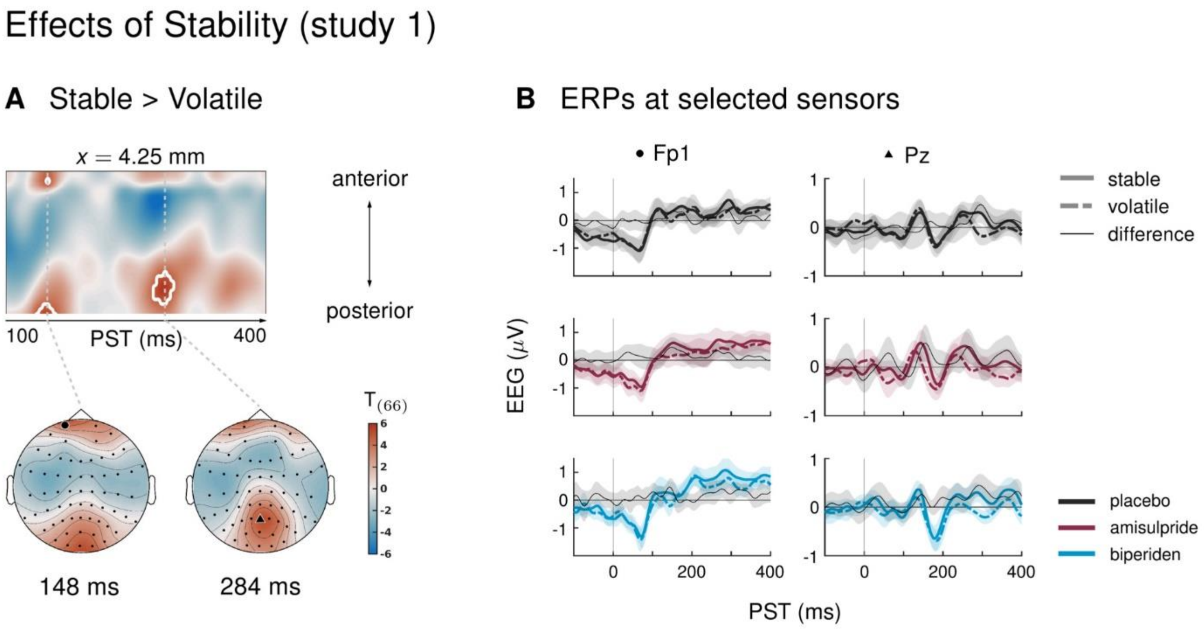


Figure S1 Main effect of stability in study 1. **A** Regions of the time × sensor space where ERPs to tones in stable phases were more positive than ERPs to tones in volatile phases. Logic of display as in Figure 2 in the main text. We found early (at 148ms in pre-frontal sensors and at 144ms in occipital sensors) and late effects (at 284ms in parietal sensors) of stability on the ERPs. **B** ERPs and difference waves for selected sensors, separately for the three drug groups. In all clusters, ERPs to tones in stable phases were more positive than ERPs to tones in volatile phases.

#### Study 1: Stability × drug group interaction

While the late effect of stability was quite consistent across drug groups, the earlier effects, particularly at pre-frontal channels, were expressed most prominently in the placebo group. In contrast, the ERPs in the biperiden group displayed a different effect early on: responses to both standard and deviant tones around 124ms at central sensors were less negative during volatile phases than during stable phases. In other words, the N1 component of the auditory ERP to tones was less pronounced in volatile phases than during stable phases, and this N1 decrement with volatility was not observed in the other groups.

However, when directly contrasting the impact of stability on the auditory ERPs between drug groups, we found no significant differences, both across the whole time × sensor space and within the functionally defined mask.

| **Effects of stability** | cluster | $x[mm]$ | $y[mm]$ | $z[ms]$ | $t_{66/73}$ | $Z_{\equiv}$ | $p_{FWE}$ | $k_E$ | $tw_{sig}$ [ms] |
|---|---|---|---|---|---|---|---|---|---|
| **A** study 1: stable > volatile | 1 | 4 | -62 | 284 | 5.32 | 4.83 | 0.003 | 110 | 272 - 292 |
| | 2 | 4 | -95 | 144 | 5.04 | 4.62 | 0.007 | 45 | 136 - 156 |
| | 3 | 0 | 61 | 148 | 4.76 | 4.39 | 0.017 | 14 | 144 - 148 |
| **B** study 2: volatile > stable | 1 | 0 | 40 | 268 | 4.51 | 4.22 | 0.035 | 6 | 268 - 272 |
| | 2 | 4 | 24 | 384 | 4.42 | 4.15 | 0.046 | 4 | 384 - 384 |
| **C** study 2: PLA > LEV | 1 | -38 | -14 | 264 | 4.58 | 4.28 | 0.028 | 4 | 264 - 264 |
| **D** study 2: GAL > PLA | 1 | -30 | 61 | 204 | 4.73 | 4.40 | 0.018 | 7 | 200 - 208 |

Table S2: Significant clusters for main effects of stability and interactions stability × drug group across the two studies. Columns are organized as in Table 1 of the main text.

#### Study 2: Main effect of stability

In study 2, ERPs to tones in volatile phases were significantly more positive than ERPs to tones in stable phases between 268ms and 272ms at frontal sensors (peak at 268ms, $t$=4.22, $p$=0.035; and at 384ms, $t$=4.42, $p$=0.046; Table S2, Figure S2A). Plotting stable and volatile ERPs at these sensors separately for the three drug groups shows a sustained difference between the conditions in the galantamine group, whereas the effects appear to be more transient in the other groups (Figure S2B).

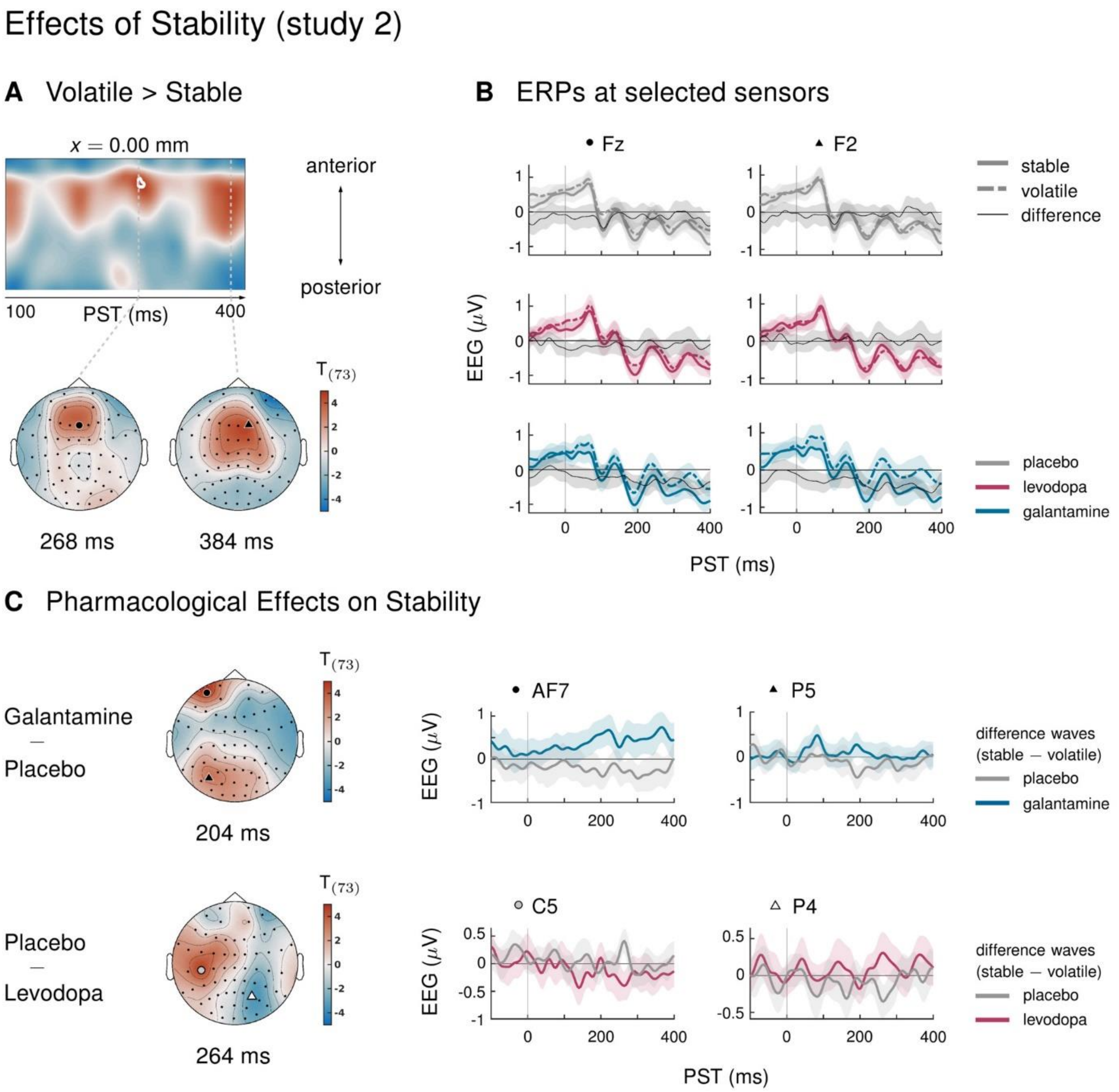


Figure S2: Main effect of stability and interaction stability × drug group in study 2. **A** Regions of the time × sensor space where ERPs to tones in volatile phases were more positive than ERPs to tones in stable phases. Logic of display as in Figure 2 in the main text. **B** ERPs and difference waves for selected sensors, separately for the three drug groups. In this cluster, ERPs to tones in volatile phases were more positive than ERPs to tones in stable phases. This effect was driven by the galantamine group. **C** Pharmacological effects on stability: we found significant differences in the impact of stability on ERPs in two clusters. Plots show the ERP difference waves (ERPs to tones in stable phases − ERPs to tones in volatile phases) in different drug conditions.

#### Study 2: Stability × drug group interaction

The impact of stability on auditory ERPs differed significantly between the drug groups in two places. Between 200ms and 208ms, left prefrontal sensors showed significantly more positive modulation by stability (ERP amplitudes being more positive for tones in stable phases) in the galantamine group compared to the placebo group (peak at 204ms, $t$=4.73, $p$=0.018). At 264ms, the levodopa group differed from the placebo group ($t$=4.58, $p$=0.028) in that left central and fronto-central sensors showed more positive amplitudes in response to tones in volatile compared to stable phases, while the opposite effect was visible in the placebo group (Figure S2C). There were no additional differences between drug groups when constraining the search volume using the average effect of stability.

#### Effects of stability: conclusions

In summary, we also found that stability itself modulated auditory ERPs, irrespective of predictability (standards vs. deviants), in both studies, although these effects appear small and only partly replicate across the two studies. The most consistent finding is an effect on later ERP components around 272ms, which manifested as a positive modulation (stable>volatile) at parietal sensors (study 1), and a negative modulation (volatile>stable) at frontal sensors (study 2). In study 2, this effect was strongly driven by the galantamine group. While an effect of galantamine on the modulation of evoked responses by stability/volatility would be intriguing, the observed effects in the present study appear as ERP-wide differences rather than modulations of specific ERP components. Our pre-processing choices (weak high-pass filtering, no baseline correction) do not exclude the possibility that slow drifts were affecting the offset of the grand averages in the stable and volatile conditions differentially. Unlike deviance, stability was manipulated in blocks rather than on a trial-by-trial basis. This might have led to the apparent ERP differences in the galantamine group.

Importantly, interaction effects between deviance and stability, as we report them in the main text, do not suffer from this constraint: the quickly alternating standard and deviant trials would be affected equally by tonic offsets, thus, any interaction of the stability effects with mismatch (standards vs. deviants) cannot be explained by the impact of artifactual slow drifts.

#### 2.2 Control analyses: Excluding data sets based on the distraction task

| **Study 1:** $N = 67$ | cluster | $x[mm]$ | $y[mm]$ | $z[ms]$ | $t_{62}$ | $Z_{\equiv}$ | $p_{FWE}$ | $k_E$ | $tw_{sig}$ [ms] |
|---|---|---|---|---|---|---|---|---|---|
| **A** standards > deviants | 1 | -13 | 2 | 172 | 15.73 | Inf | 0.000 | 8060 | 100 - 228 |
| | | 8 | -3 | 180 | 15.26 | Inf | 0.000 | | |
| | 2 | 4 | 2 | 400 | 9.17 | 7.26 | 0.000 | 1181 | 364 - 400 |
| | 3 | -8 | 50 | 276 | 6.80 | 5.85 | 0.000 | 264 | 240 - 300 |
| | | -4 | 45 | 248 | 5.03 | 4.59 | 0.007 | | |
| | 4 | 0 | -95 | 304 | 5.13 | 4.66 | 0.005 | 92 | 288 - 332 |
| | 5 | -60 | -57 | 268 | 4.83 | 4.44 | 0.013 | 191 | 240 - 284 |
| | | -60 | -9 | 256 | 4.75 | 4.37 | 0.016 | | |
| | | -60 | -36 | 260 | 4.74 | 4.37 | 0.017 | | |
| **B** deviants > standards | 1 | 17 | 72 | 156 | 13.99 | Inf | 0.000 | 1524 | 100 - 236 |
| | | 4 | 72 | 212 | 9.74 | 7.56 | 0.000 | | |
| | | -30 | 61 | 184 | 7.67 | 6.41 | 0.000 | | |
| | 2 | -47 | -68 | 180 | 13.96 | Inf | 0.000 | 5986 | 100 - 328 |
| | | 64 | -62 | 200 | 13.52 | Inf | 0.000 | | |
| | | 42 | -78 | 168 | 11.49 | Inf | 0.000 | | |
| | 3 | -51 | -30 | 400 | 6.18 | 5.44 | 0.000 | 340 | 372 - 400 |
| | 4 | 34 | -52 | 364 | 6.12 | 5.39 | 0.000 | 418 | 352 - 400 |
| | | 47 | -52 | 400 | 5.60 | 5.02 | 0.001 | | |
| | 5 | 64 | -62 | 100 | 4.87 | 4.46 | 0.012 | 9 | 100 - 112 |
| | 6 | 0 | 72 | 400 | 4.67 | 4.31 | 0.021 | 8 | 400 - 400 |
| | | -17 | 72 | 400 | 4.49 | 4.17 | 0.035 | | |
| **C** mismatch: AMI > BIP | 1 | 13 | 67 | 160 | 5.26 | 4.76 | 0.003 | 70 | 148 - 168 |
| **D** stable MMN > volatile MMN | 1 | 17 | -19 | 204 | 5.25 | 4.75 | 0.004 | 407 | 180 - 216 |
| | | -17 | -25 | 196 | 5.20 | 4.72 | 0.005 | | |
| | 2 | -8 | 45 | 256 | 4.70 | 4.33 | 0.022 | 19 | 252 - 260 |
| **E** volatile MMN > stable MMN | 1 | 42 | -78 | 196 | 5.12 | 4.66 | 0.006 | 69 | 188 - 212 |
| | | 30 | -89 | 204 | 4.89 | 4.48 | 0.013 | | |
| | | 55 | -68 | 196 | 4.83 | 4.43 | 0.015 | | |
| | 2 | -60 | -57 | 208 | 5.00 | 4.56 | 0.009 | 15 | 200 - 216 |
| | 3 | 4 | 72 | 212 | 4.54 | 4.20 | 0.036 | 7 | 208 - 212 |
| | 4 | 13 | -95 | 208 | 4.49 | 4.16 | 0.041 | 2 | 208 - 212 |
| **F** stable > volatile | 1 | 4 | -62 | 284 | 5.54 | 4.98 | 0.001 | 156 | 272 - 296 |
| | 2 | 4 | -95 | 144 | 4.88 | 4.47 | 0.013 | 35 | 140 - 152 |
| | 3 | 0 | 61 | 148 | 4.50 | 4.17 | 0.039 | 2 | 144 - 148 |

Table S3 All clusters of significant activation for a reduced sample size (*N*=67) in study 1. Table shows whole-volume corrected significant effects after excluding three data sets due to lack of behavioral data and one dataset based on low performance in the distraction task (hit rate<75%). All results reported for the full sample (*N*=71, Tables 1, 2, S1) hold.

| **Study 2:** $N = 77$ | cluster | $x[mm]$ | $y[mm]$ | $z[ms]$ | $t_{72}$ | $Z_{\equiv}$ | $p_{FWE}$ | $k_E$ | $tw_{sig}$ [ms] |
|---|---|---|---|---|---|---|---|---|---|
| **A** standards > deviants | 1 | 13 | -9 | 176 | 13.99 | Inf | 0.000 | 7462 | 100 - 216 |
| | | 4 | 18 | 160 | 13.79 | Inf | 0.000 | | |
| | | 42 | -19 | 124 | 10.61 | Inf | 0.000 | | |
| | 2 | 0 | -3 | 396 | 8.36 | 6.95 | 0.000 | 1317 | 364 - 400 |
| | 3 | 4 | -95 | 296 | 7.25 | 6.25 | 0.000 | 688 | 244 - 332 |
| | | 30 | -89 | 280 | 6.64 | 5.83 | 0.000 | | |
| | | 51 | -68 | 256 | 5.53 | 5.02 | 0.001 | | |
| | 4 | 4 | 61 | 288 | 6.07 | 5.43 | 0.000 | 290 | 256 - 304 |
| | | 0 | 56 | 268 | 5.92 | 5.32 | 0.000 | | |
| | 5 | -47 | -68 | 256 | 5.78 | 5.21 | 0.000 | 298 | 232 - 284 |
| | | -60 | -57 | 260 | 5.61 | 5.09 | 0.001 | | |
| **B** deviants > standards | 1 | -42 | -73 | 172 | 14.11 | Inf | 0.000 | 5568 | 100 - 216 |
| | | 55 | -68 | 196 | 10.81 | Inf | 0.000 | | |
| | | -38 | -73 | 124 | 10.15 | Inf | 0.000 | | |
| | 2 | -13 | -30 | 256 | 7.31 | 6.29 | 0.000 | 2724 | 232 - 328 |
| | | -26 | -14 | 288 | 6.85 | 5.98 | 0.000 | | |
| | | 8 | -9 | 304 | 6.61 | 5.81 | 0.000 | | |
| | 3 | 38 | -68 | 400 | 6.46 | 5.71 | 0.000 | 273 | 368 - 400 |
| | 4 | 4 | 72 | 388 | 5.86 | 5.28 | 0.000 | 146 | 368 - 400 |
| | 5 | 68 | 18 | 196 | 5.39 | 4.92 | 0.002 | 16 | 172 - 204 |
| | 6 | -60 | -9 | 400 | 4.95 | 4.57 | 0.008 | 36 | 396 - 400 |
| | 7 | -34 | -62 | 396 | 4.86 | 4.50 | 0.010 | 50 | 388 - 400 |
| **C** volatile > stable | 1 | 4 | 18 | 384 | 4.63 | 4.32 | 0.025 | 32 | 380 - 388 |
| | 2 | 0 | 40 | 268 | 4.47 | 4.18 | 0.041 | 3 | 268 - 272 |
| | 3 | -21 | -19 | 356 | 4.41 | 4.13 | 0.050 | 1 | 356 - 356 |
| **D** stability: PLA > LEV | 1 | -38 | -14 | 264 | 4.70 | 4.37 | 0.020 | 9 | 260 - 264 |

Table S4 All clusters of significant activation for a reduced sample size (*N*=77) in study 2. Table shows whole-volume corrected significant effects after excluding one data set based on low performance in the distraction task (hit rate<75%). All results reported for the full sample (*N*=78, Tables 3, S2) hold, except for the pharmacological effect (GAL > PLA) on stability ERPs reported in Table S2.

#### 2.3 Genetic effects and pharmaco-genetic interactions

To examine effects of individual differences in dopamine metabolism on mismatch related ERPs and potential interactions of these differences with the effects of amisulpride, we added a covariate coding for genetic variations (SNPs) in the COMT polymorphism to the group level GLMs (see Supplementary Methods section 1.3). In study 1, we found a significant modulation of mismatch ERPs by COMT SNP in the amisulpride group in parieto-occipital sensors (Val/Val > Val/Met > Met/Met, 300-308ms, peak at 304ms, $t$=4.51, $p$=0.033, Table S5), and in right central sensors (Met/Met > Val/Met > Val/Val, 292-304ms, peak at 300ms, $t$=4.79, $p$=0.014, Table S5).

| **Study 1: Genetic effects** | cluster | $x[mm]$ | $y[mm]$ | $z[ms]$ | $t_{63}$ | $Z_{\equiv}$ | $p_{FWE}$ | $k_E$ | $tw_{sig}$ [ms] |
|---|---|---|---|---|---|---|---|---|---|
| **A** volatile mismatch: CHAT: PLA > BIP | 1 | -34 | -52 | 380 | 5.77 | 5.15 | 0.001 | 302 | 356 - 392 |
| **B** volatile mismatch: PLA: pos. effect of CHAT | 1 | -34 | -46 | 384 | 4.57 | 4.23 | 0.032 | 6 | 380 - 384 |
| **C** volatile mismatch: BIP: neg. effect of CHAT | 1 | -51 | -25 | 384 | 4.49 | 4.17 | 0.040 | 6 | 380 - 384 |
| **D** mismatch: AMI: pos. effect of COMT | 1 | 0 | -73 | 304 | 4.51 | 4.18 | 0.033 | 18 | 300 - 308 |
| **E** mismatch: AMI: neg. effect of COMT | 1 | 42 | -19 | 300 | 4.79 | 4.41 | 0.014 | 33 | 292 - 304 |
| **F** stable mismatch: AMI: neg. effect of COMT | 1 | 42 | -14 | 300 | 4.44 | 4.12 | 0.042 | 6 | 300 - 304 |

Table S5 Genetic effects and pharmaco-genetic interactions in study 1. Columns are organized as in Table 1 of the main text. Mismatch ERPs in volatile periods were differentially modulated by CHAT polymorphism in the placebo versus the biperiden group, whereas mismatch ERPs overall and in stable periods were modulated by COMT polymorphism in the amisulpride group.

Similarly, in separate GLMs, we considered effects of the CHAT polymorphism on mismatch related ERPs in the placebo and the biperiden group and potential interactions of CHAT SNPs with the effect of biperiden. Here, we found a significant modulation of mismatch ERPs in volatile periods of the experiment by CHAT polymorphism. At left centro-parietal sensors at 384ms after tone onset, volatile mismatch effects depended on CHAT SNP within the placebo group (A/A > A/G > G/G, $t=4.57$, $p=0.032$, Table S5) and this modulation was significantly different from the effect of CHAT in the biperiden group at these sensors (significant interaction between CHAT covariate and drug group, 356-492ms, $t=5.77$, $p=0.001$, Table S5).

| **Study 2: Genetic effects** | cluster | $x[mm]$ | $y[mm]$ | $z[ms]$ | $t_{68}$ | $Z_{\equiv}$ | $p_{FWE}$ | $k_E$ | $tw_{sig}$ [ms] |
|---|---|---|---|---|---|---|---|---|---|
| **A** volatile mismatch: PLA: pos. effect of CHAT | 1 | -13 | -78 | 104 | 4.42 | 4.13 | 0.046 | 4 | 104 - 108 |
| **B** volatile mismatch: PLA: neg. effect of COMT | 1 | 4 | 45 | 192 | 5.31 | 4.84 | 0.003 | 53 | 180 - 200 |

Table S6 Genetic effects and pharmaco-genetic interactions in study 2. Columns are organized as in Table 1 of the main text. Volatile mismatch ERPs in the placebo group were positively modulated by CHAT polymorphism, and negatively modulated by COMT polymorphism.

In study 2, we found that the amplitude of mismatch ERPs in volatile periods of the experiment depended both on CHAT and COMT polymorphism, but only in the placebo group (see Table S6 for details).

However, all of these results (modulation of mismatch ERPs by COMT in the amisulpride group and differential modulation of volatile mismatch ERPs by CHAT in the placebo compared to the biperiden group, modulation of volatile mismatch ERPs by both CHAT and COMT in the placebo group of study 2) have to be treated with great caution, given that our sample size is very small for detecting (the presumably subtle) genetic effects. For example, in study 1, for COMT, there were only 3 individuals showing the Val/Val genotype in the amisulpride group, and for CHAT, there were only 2 individuals showing the A/A genotype in the biperiden group, and 3 individuals with this genotype in the placebo group. Moreover, none of the genetic effects in the placebo groups replicated across the two studies. Given these limitations, we refrain from interpreting or discussing these genetic effects any further and report them here only for completeness and transparency, and as potential guidance for future follow-up studies with larger sample sizes.

#### 2.4 Robustness of our results against analysis choices

##### Alternative pre-processing and trial definition

We re-examined our main results under a different pre-processing strategy (including baseline correction, a strong high-pass filter, and a linked mastoid reference), and a different trial definition (retaining more trials per condition). Additionally, we complemented our whole time × sensor analysis with a region-of-interest (ROI) analysis based on previous literature to maximize our sensitivity to drug effects. We were particularly interested in *(1)* the main effect of mismatch and its modulation by drug group, and *(2)* the interaction of mismatch and stability after correcting for slow drifts and increasing trial numbers.

*1) Pharmacological effects on mismatch*

Despite a very different pre-processing strategy and a different trial definition, the main effects of mismatch in both studies were highly similar to what we reported in the main text: a typical fronto-central MMN cluster between 100 and 244ms, followed by a fronto-central P3a like response between 232 and 352ms, and a late P3b-like effect between 376 and 400ms (with all fronto-central effects being mirrored by opposite-sign effects in prefrontal and temporal sensors, as is typical for the MMN, see Tables S7,S8, Figure S3A,B).

The new analysis located the most prominent pharmacological effect in a later time window compared to the results with our original pipeline: The late positive component of the mismatch waveform showed a delayed peak under biperiden, resulting in significant differences in mismatch amplitude between the biperiden group and both the amisulpride and the placebo group around 344ms (Table S7, Figure S3C). Such a shift in the dominant ERP component is not surprising when using a strong high-pass filter (for a critical discussion of the effects of strong high-pass filtering, see (Tanner et al., 2015)).

| **Study 1: mismatch** | cluster | $x[mm]$ | $y[mm]$ | $z[ms]$ | $t_{66}$ | $Z_{\equiv}$ | $p_{FWE}$ | $k_E$ | $tw_{sig}$ [ms] |
|---|---|---|---|---|---|---|---|---|---|
| **A** standards > deviants | | -8 | 8 | 176 | 16.16 | Inf | 0.000 | 15744 | 100 - 244 |
| | | 13 | 8 | 172 | 16.06 | Inf | 0.000 | | |
| | | 51 | -25 | 140 | 15.26 | Inf | 0.000 | | |
| | | 8 | 56 | 284 | 9.53 | 7.57 | 0.000 | 2142 | 244 - 364 |
| | | 0 | -9 | 400 | 5.91 | 5.29 | 0.000 | 533 | 384 - 400 |
| **B** deviants > standards | | 13 | 72 | 168 | 16.55 | Inf | 0.000 | 2501 | 100 - 220 |
| | | -17 | -9 | 280 | 8.79 | 7.16 | 0.000 | 9514 | 244 - 376 |
| | | 0 | 2 | 308 | 8.44 | 6.96 | 0.000 | | |
| | | 42 | -3 | 324 | 7.81 | 6.57 | 0.000 | | |
| | | 38 | 56 | 400 | 5.22 | 4.77 | 0.002 | 55 | 388 - 400 |
| | | -60 | -57 | 132 | 4.96 | 4.57 | 0.005 | 13 | 116 - 144 |
| | | -42 | 45 | 400 | 4.62 | 4.29 | 0.014 | 18 | 396 - 400 |
| **C** BIP > PLA | | 8 | -14 | 344 | 4.87 | 4.50 | 0.006 | 144 | 336 - 352 |
| | | -26 | 45 | 344 | 5.06 | 4.64 | 0.003 | 104 | 332 - 352 |
| **D** BIP > AMI | | -21 | 45 | 344 | 4.88 | 4.50 | 0.006 | 44 | 340 - 356 |

Table S7 Main effects of mismatch and interaction mismatch*drug in study 1 under the new pre-processing pipeline and trial definition. Columns are organized as in Table 1 of the main text. Mismatch responses under biperiden were significantly different from placebo and amisulpride around 344ms.

| **Study 2: mismatch** | cluster | $x[mm]$ | $y[mm]$ | $z[ms]$ | $t_{73}$ | $Z_{\equiv}$ | $p_{FWE}$ | $k_E$ | $tw_{sig}$ [ms] |
|---|---|---|---|---|---|---|---|---|---|
| **A** standards > deviants | | -13 | 18 | 156 | 14.88 | Inf | 0.000 | 14562 | 100 - 244 |
| | | 51 | -14 | 136 | 13.55 | Inf | 0.000 | | |
| | | 0 | 72 | 296 | 11.89 | Inf | 0.000 | 2425 | 236 - 352 |
| | | 4 | 2 | 400 | 9.45 | 7.61 | 0.000 | 1084 | 376 - 400 |
| **B** deviants > standards | | 0 | 72 | 156 | 14.14 | Inf | 0.000 | 2260 | 100 - 208 |
| | | 34 | 8 | 308 | 12.49 | Inf | 0.000 | 14460 | 232 - 400 |
| | | 38 | 2 | 284 | 12.19 | Inf | 0.000 | | |
| | | -26 | 2 | 296 | 11.59 | Inf | 0.000 | | |

Table S8 Main effects of mismatch in study 2 under the new pre-processing pipeline and trial definition. Columns are organized as in Table 1 of the main text. There were no clusters with significant differences between drug groups.

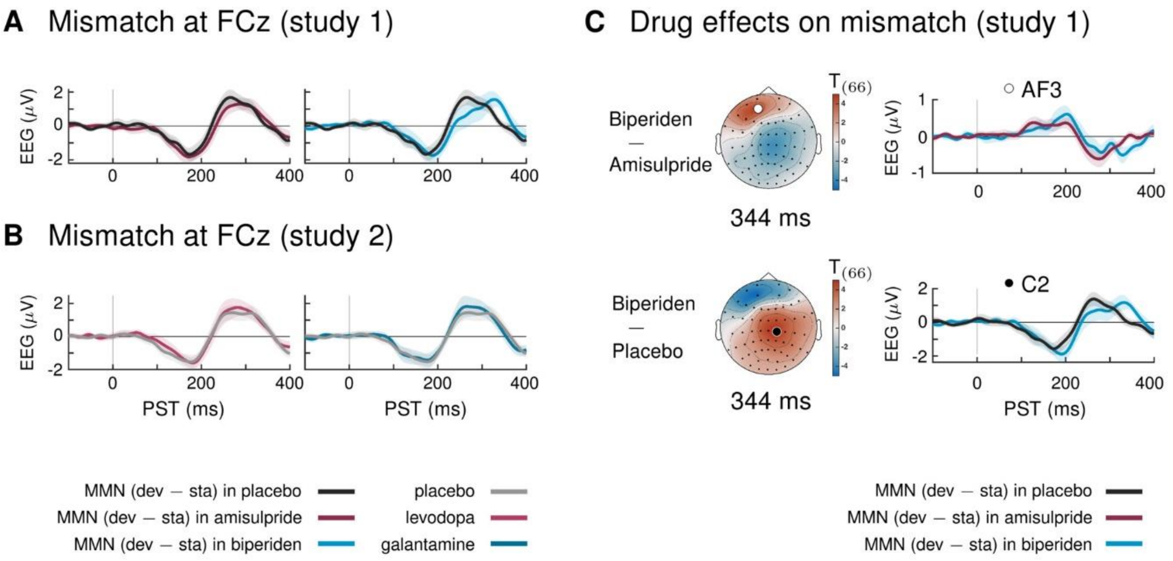


Figure S3: Main effect of mismatch and interaction mismatch × drug group under the new pre-processing pipeline and trial definition. **A** Mismatch signals (deviants – standards) at sensor FCz in study 1. **B** Mismatch signals (deviants – standards) at sensor FCz in study 2. There were no significant differences between mismatch signals of the different drug groups in study 2. **C** Pharmacological effects on mismatch in study 1: the new pipeline located the dominant effect of biperiden in a later time window compared to our original pipeline. Around 344ms, mismatch signals were significantly stronger in the biperiden group compared to placebo and amisulpride, due to a delayed peak of the P3 component.

Critically, and consistent with our findings in the main text, mismatch responses were affected by biperiden, compared to placebo and amisulpride, while we did not find any evidence for dopaminergic effects on mismatch responses. Just as in our main analysis, there were no significant effects of drug on mismatch responses in study 2 (galantamine and levodopa).

To further ascertain that our effects are comparable to previous reports on muscarinergic effects on MMN in the literature (Klinkenberg et al., 2013; Caldenhove et al., 2017) and that our lack of galantamine effects on MMN (in contrast to (Moran et al., 2013)) was not due to low sensitivity of our whole time × sensor analysis, we additionally performed a region-of-interest (ROI) analysis, focusing on exactly those sensors used in these previous studies, and following their approach for extracting MMN amplitudes and latencies in every participant (Klinkenberg et al., 2013; Caldenhove et al., 2017).

In line with the results reported in the main text, we found that the MMN peak latency in study 1 was increased under biperiden (mean: 181.9ms, std: 3.4) compared to placebo (mean: 168.8ms, std: 3.2, Table S9). Peak amplitudes were not significantly different between drug groups. This is consistent with a temporal shift of the mismatch response in the early (classical) MMN time window of the kind we describe in the main text. In other words, even though the whole time × sensor space analysis under the new processing pipeline had located the dominant drug effect in a later component, we still find evidence for this early MMN shift when focusing on the classical MMN sensors, supporting the original results reported in the main manuscript.

| | Study 1 | | | | Study 2 | | | |
|---|---|---|---|---|---|---|---|---|
| | PLA | AMI | BIP | $F_{2,208}$ (*p*) | PLA | LEV | GAL | *F* (*p*) |
| **Peaks (µV)** | -1.83 (0.1) | -1.95 (0.1) | -2.04 (0.1) | 1.1 *(0.33)* | -1.75 (0.1) | -2.0 (0.1) | -1.76 (0.1) | 1.73 (0.18) |
| **Lat. (ms)** | 168.8 (3.2) | 173.7 (3.2) | 181.9 (3.4) | **4.06** ***(0.02)*** | 165.1 (2.75) | 173.2 (2.75) | 163.7 (2.75) | **3.5** ***(0.03)*** |
| Post-hoc *t* | Lat.: BIP>PLA | | ***p=0.013*** | | Lat.: LEV>GAL | | ***p=0.038*** | |

Table S9 Results of the ROI analysis. Table lists mean (std) values of the peak amplitudes and latencies separately for each drug group in the two studies. *F* (*p*) values refer to the effect of the factor drug group in a 3x3 ANOVA (drug x sensor). Last row lists the significant post-hoc comparisons between pairs of drug groups. Lat. = Latency.

In study 2, we again found no significant differences in MMN peak amplitude between drug groups. However, peak latency was significantly different between the galantamine and the levodopa group, such that mismatch responses peaked earlier under galantamine (Table S9). This suggests that while blocking (muscarinic) cholinergic receptors delays MMN, enhancing cholinergic processing via galantamine might result in faster mismatch responses. However, peak latency under galantamine was not significantly different from placebo. Moreover, we did not observe any effect of galantamine in any other analysis, suggesting that this effect was subtle at most.

*2) Stability effects on mismatch processing*

Supporting our claim of stronger mismatch signals during stable phases of the experiment, we found a significant interaction between mismatch and stability irrespective of the pre-processing pipeline and the trial definition (see Tables S10,11 and Figure S4). In particular, in all cases (original and new pre-processing, original and new trial definition), mismatch signals were significantly stronger during stable phases of the experiment in left central and fronto-central sensors around 200ms (Tables S10,11, Figure S4). The only exception to this was study 2 when analyzed with our original pre-processing pipeline (no baseline correction, weak high-pass filter) in combination with our original trial definition (which resulted in a relatively low number of trials per condition). As reported in the main text, under this pipeline, no significant interaction effect emerged. This suggests that for examining the effects of block-wise changes in stability on mismatch processing, a more sensitive analysis approach (by removing slow drifts or increasing trial numbers) is needed. It is also consistent with a more subtle interaction effect in study 2 compared to study 1, presumably because the interaction effect was particularly pronounced in the biperiden group in study 1. Importantly, the interaction effect in study 1 as reported in the main text was robust against analysis choices and is thus not an artefact of the specific processing steps we applied.

| **Mismatch × stability: study 1** | cluster | $x[mm]$ | $y[mm]$ | $z[ms]$ | $t_{66}$ | $Z_{\equiv}$ | $p_{FWE}$ | $k_E$ | $tw_{sig}$ [ms] |
|---|---|---|---|---|---|---|---|---|---|
| **A** ORG preprocessing, ROV trial definition | | | | | | | | | |
| stable > volatile mismatch (pos.) | 1 | -21 | -25 | 196 | 5.69 | 5.11 | 0.001 | 581 | 180 - 220 |
| | | 17 | -19 | 208 | 5.68 | 5.11 | 0.001 | | |
| stable > volatile mismatch (neg.) | 1 | 42 | -78 | 200 | 5.70 | 5.12 | 0.001 | 118 | 188 - 232 |
| | | 55 | -68 | 200 | 5.21 | 4.75 | 0.004 | | |
| | | 64 | -62 | 200 | 5.06 | 4.64 | 0.006 | | |
| | 2 | -60 | -57 | 208 | 5.23 | 4.77 | 0.004 | 22 | 204 - 220 |
| | 3 | -60 | -57 | 124 | 4.53 | 4.21 | 0.034 | 2 | 120 - 124 |
| **B** ORG preprocessing, FAIR trial definition | | | | | | | | | |
| stable > volatile mismatch (pos.) | 1 | -13 | -19 | 176 | 9.68 | 7.61 | 0.000 | 3786 | 136 - 256 |
| | | -8 | -9 | 208 | 7.99 | 6.66 | 0.000 | | |
| | | 17 | -9 | 216 | 7.56 | 6.39 | 0.000 | | |
| stable > volatile mismatch (neg.) | 1 | -42 | -73 | 216 | 6.66 | 5.80 | 0.000 | 977 | 144 - 236 |
| | | 30 | -89 | 168 | 6.57 | 5.74 | 0.000 | | |
| | | -26 | -89 | 216 | 6.38 | 5.61 | 0.000 | | |
| | 2 | 4 | 67 | 212 | 5.68 | 5.10 | 0.001 | 510 | 152 - 236 |
| | | 0 | 61 | 168 | 5.62 | 5.06 | 0.001 | | |
| | | -21 | 67 | 224 | 5.27 | 4.80 | 0.003 | | |
| | 3 | -42 | -73 | 140 | 4.46 | 4.15 | 0.039 | 2 | 136 - 140 |
| **C** NEW preprocessing, ROV trial definition | | | | | | | | | |
| stable > volatile mismatch (pos.) | 1 | -21 | -9 | 208 | 5.38 | 4.90 | 0.001 | 1083 | 192 - 240 |
| | | -26 | -30 | 224 | 5.25 | 4.79 | 0.002 | | |
| | | 21 | -19 | 212 | 4.62 | 4.30 | 0.017 | | |
| stable > volatile mismatch (neg.) | 1 | 4 | 56 | 216 | 5.71 | 5.14 | 0.000 | 331 | 200 - 232 |
| | | 13 | 61 | 208 | 5.66 | 5.10 | 0.001 | | |
| | 2 | -4 | 50 | 120 | 5.27 | 4.81 | 0.002 | 36 | 112 - 128 |
| **D** NEW preprocessing, FAIR trial definition | | | | | | | | | |
| stable > volatile mismatch (pos.) | 1 | -30 | -14 | 176 | 6.17 | 5.48 | 0.000 | 3330 | 140 - 232 |
| | | 26 | -9 | 216 | 5.98 | 5.34 | 0.000 | | |
| | | -13 | -14 | 208 | 5.84 | 5.24 | 0.000 | | |
| | 2 | -26 | 45 | 360 | 4.53 | 4.22 | 0.023 | 5 | 360 - 364 |
| stable > volatile mismatch (neg.) | 1 | -4 | 50 | 192 | 6.84 | 5.94 | 0.000 | 879 | 164 - 228 |
| | | 4 | 56 | 216 | 6.45 | 5.68 | 0.000 | | |
| | 2 | 38 | -52 | 364 | 4.82 | 4.45 | 0.009 | 127 | 360 - 372 |
| | 3 | 0 | 45 | 128 | 4.78 | 4.42 | 0.011 | 27 | 120 - 136 |

Table S10 Interaction effects mismatch*stability in study 2 under the different pre-processing pipelines and trial definitions. Columns are organized as in Table 1 of the main text. In all cases, mismatch was larger (higher positivity in prefrontal and temporal sensors; higher negativity in frontocentral sensors) during stable phases of the experiment before and around 200ms.

| **Mismatch × stability: study 2** | cluster | $x[mm]$ | $y[mm]$ | $z[ms]$ | $t_{73}$ | $Z_{\equiv}$ | $p_{FWE}$ | $k_E$ | $tw_{sig}$ [ms] |
|---|---|---|---|---|---|---|---|---|---|
| **A** ORG preprocessing, ROV trial definition | | | | | | | | | |
| *no sig. clusters* | | | | | | | | | |
| **B** ORG preprocessing, FAIR trial definition | | | | | | | | | |
| stable > volatile mismatch (pos.) | 1 | -17 | 29 | 224 | 6.50 | 5.75 | 0.000 | 2122 | 172 - 284 |
| | | -13 | 45 | 252 | 6.39 | 5.68 | 0.000 | | |
| | | -21 | 8 | 184 | 6.26 | 5.58 | 0.000 | | |
| | 2 | -13 | 40 | 400 | 5.49 | 5.01 | 0.001 | 101 | 356 - 400 |
| | | -4 | 45 | 368 | 4.88 | 4.53 | 0.011 | | |
| | 3 | 21 | -3 | 120 | 4.48 | 4.20 | 0.039 | 10 | 116 - 120 |
| | 4 | -21 | 29 | 144 | 4.40 | 4.13 | 0.049 | 1 | 144 - 144 |
| stable > volatile mismatch (neg.) | 1 | -47 | -68 | 172 | 6.53 | 5.77 | 0.000 | 278 | 164 - 208 |
| | 2 | 55 | -68 | 188 | 5.38 | 4.92 | 0.002 | 178 | 172 - 232 |
| | | 55 | -68 | 224 | 4.60 | 4.30 | 0.027 | | |
| | 3 | 4 | 72 | 192 | 5.26 | 4.83 | 0.003 | 84 | 176 - 208 |
| | 4 | 34 | 61 | 224 | 4.70 | 4.38 | 0.020 | 13 | 216 - 228 |
| | | 26 | 67 | 220 | 4.69 | 4.37 | 0.020 | | |
| **C** NEW preprocessing, ROV trial definition | | | | | | | | | |
| stable > volatile mismatch (pos.) | 1 | -26 | -41 | 208 | 6.02 | 5.40 | 0.000 | 2639 | 176 - 232 |
| | 2 | 21 | -41 | 200 | 5.20 | 4.78 | 0.002 | | |
| | 3 | 30 | -89 | 400 | 4.60 | 4.30 | 0.017 | 30 | 396 - 400 |
| stable > volatile mismatch (neg.) | 1 | 4 | 50 | 192 | 7.39 | 6.37 | 0.000 | 699 | 180 - 228 |
| | 2 | 0 | 29 | 312 | 4.44 | 4.17 | 0.027 | 12 | 308 - 312 |
| **D** NEW preprocessing, FAIR trial definition | | | | | | | | | |
| stable > volatile mismatch (pos.) | 1 | 4 | -19 | 200 | 7.28 | 6.29 | 0.000 | 6806 | 128 - 244 |
| | | -26 | -14 | 184 | 7.12 | 6.19 | 0.000 | | |
| | | -34 | -14 | 148 | 6.96 | 6.08 | 0.000 | | |
| stable > volatile mismatch (neg.) | 1 | 0 | 56 | 188 | 8.64 | 7.15 | 0.000 | 1495 | 128 - 236 |
| | | -8 | 67 | 140 | 6.36 | 5.65 | 0.000 | | |

Table S11 Interaction effects mismatch*stability in study 2 under the different pre-processing pipelines and trial definitions. Columns are organized as in Table 1 of the main text. In all cases, except the original pipeline (ORG) and trial definition (ROV), mismatch was larger (higher positivity in prefrontal and temporal sensors; higher negativity in frontocentral sensors) during stable phases of the experiment before and around 200ms.

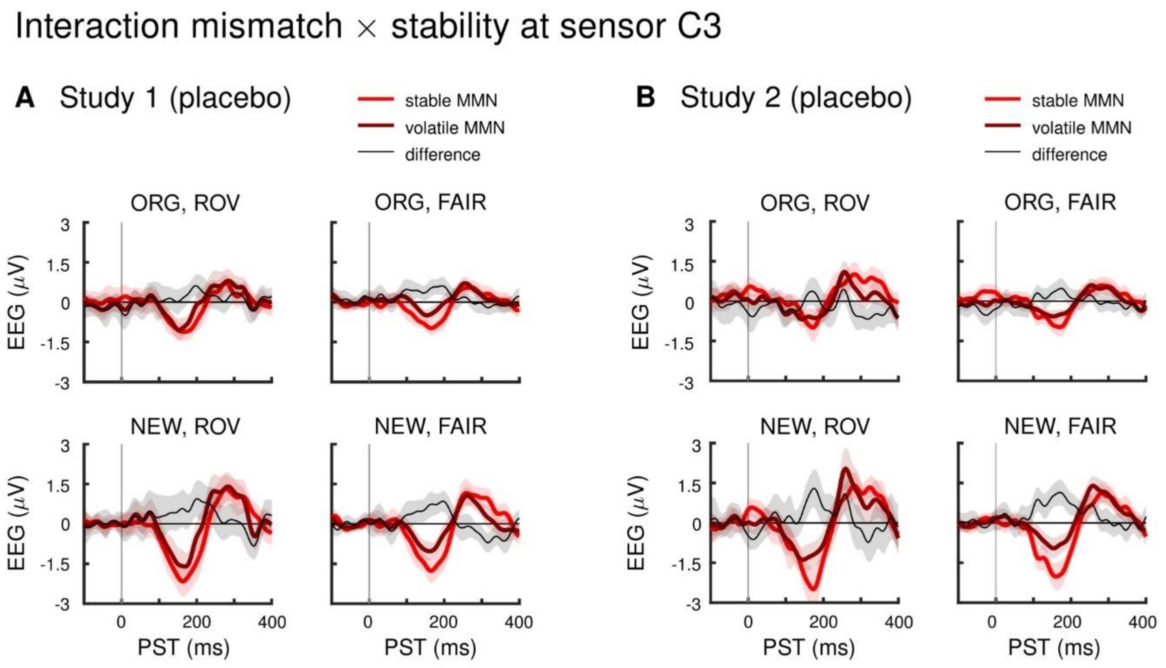


Figure S4: Interaction effect mismatch × stability at a left central sensor across pipelines. Panels show mismatch difference waves (deviants – standards) during stable and volatile phases of the experiment, and the difference between these (volatile – stable), across variations in pre-processing (ORG vs NEW) and trial definition (ROV vs FAIR). Data are shown at an exemplar left central sensor (C3; note that the location in sensor space of the peak effect varied slightly across pipelines). **A** Study 1: Stable mismatch responses were stronger than volatile mismatch responses around or before 200ms irrespective of the analysis pipeline used. **B** Study 2: Under all pipelines except ORG preprocessing and ROV trial definition, stable mismatch responses were stronger than volatile mismatch responses around or before 200ms. ORG = original pre-processing pipeline as defined in the main text, NEW = adjusted pipeline as defined in section 1.4 of this Supplementary Material, ROV = "roving" trial definition: all standards and deviants following at least 2 repetitions, FAIR = "fair" trial definition: all standards and deviants following at least 5 repetitions, for exact definitions refer to section "First level general linear model" of the main text and section 1.4 of the SM.

#### A model-based perspective on trial-by-trial auditory ERPs

To complement the perspective of our conventional ERP analysis in the main text, we also performed an analysis of auditory ERPs in our task based on the predictions of an ideal Bayesian observer. In this analysis, instead of averaging the EEG response to tones assigned to different categories ('standards', 'deviants'), we extracted an estimate of precision-weighted prediction error elicited by each tone in the sequence, according to a hierarchical Bayesian model of inference and learning (Mathys et al., 2011). This model provides prediction errors on two levels of a belief hierarchy: low-level prediction errors about tone occurrences, which serve to update beliefs about the current tendency in the environment to present one or the other tone, and higher-level prediction errors about tone probabilities, which serve to update an estimate of the current level of environmental volatility. In the following, we report clusters in which trial-wise EEG amplitudes varied in accordance with either of these PEs, and the effects of the pharmacological manipulations on the relationship between EEG and PEs.

##### *Low-level precision-weighted prediction error*

In both studies, and across drug groups, there was a significant relation between $\varepsilon_2$, our model-based trial-wise estimate of low-level precision-weighted PE, and trial-wise EEG activity corresponding to the classical MMN in frontal, fronto-central and central sensors, as well as some later clusters (Figures S4A,5A; Tables S12,13).

In study 1, we found a significant effect of drug group on the relationship between $\varepsilon_2$ and EEG amplitudes. Compared to the placebo group, ERP amplitudes in the biperiden group showed a weaker positive relation with $\varepsilon_2$ at left central sensors around 330ms after tone onset (Figure S4B; and see Table S12 for two additional very small clusters). There were no significant differences between drug groups in the effect of $\varepsilon_2$ in study 2.

##### *High-level precision-weighted prediction error*

In both studies, and across drug groups, we found significant trial-by-trial relations between $\varepsilon_3$ (the precision-weighted PE that serves to update volatility estimates) and EEG amplitudes mainly corresponding to a P3-like late central positivity (roughly 250-400ms, peaking around 300ms) and some additional smaller clusters (Figures S4C,5C; Tables S12,13).

In study 1, we found a significant effect of drug group on the relationship between $\varepsilon_3$ and EEG amplitudes. Compared to the placebo group, trial-wise amplitudes of the late central positivity in the biperiden group showed a stronger relation with $\varepsilon_3$ around 360ms after tone onset (Figure S4B; Table S12). There were no significant differences between drug groups in the effect of $\varepsilon_3$ in study 2.

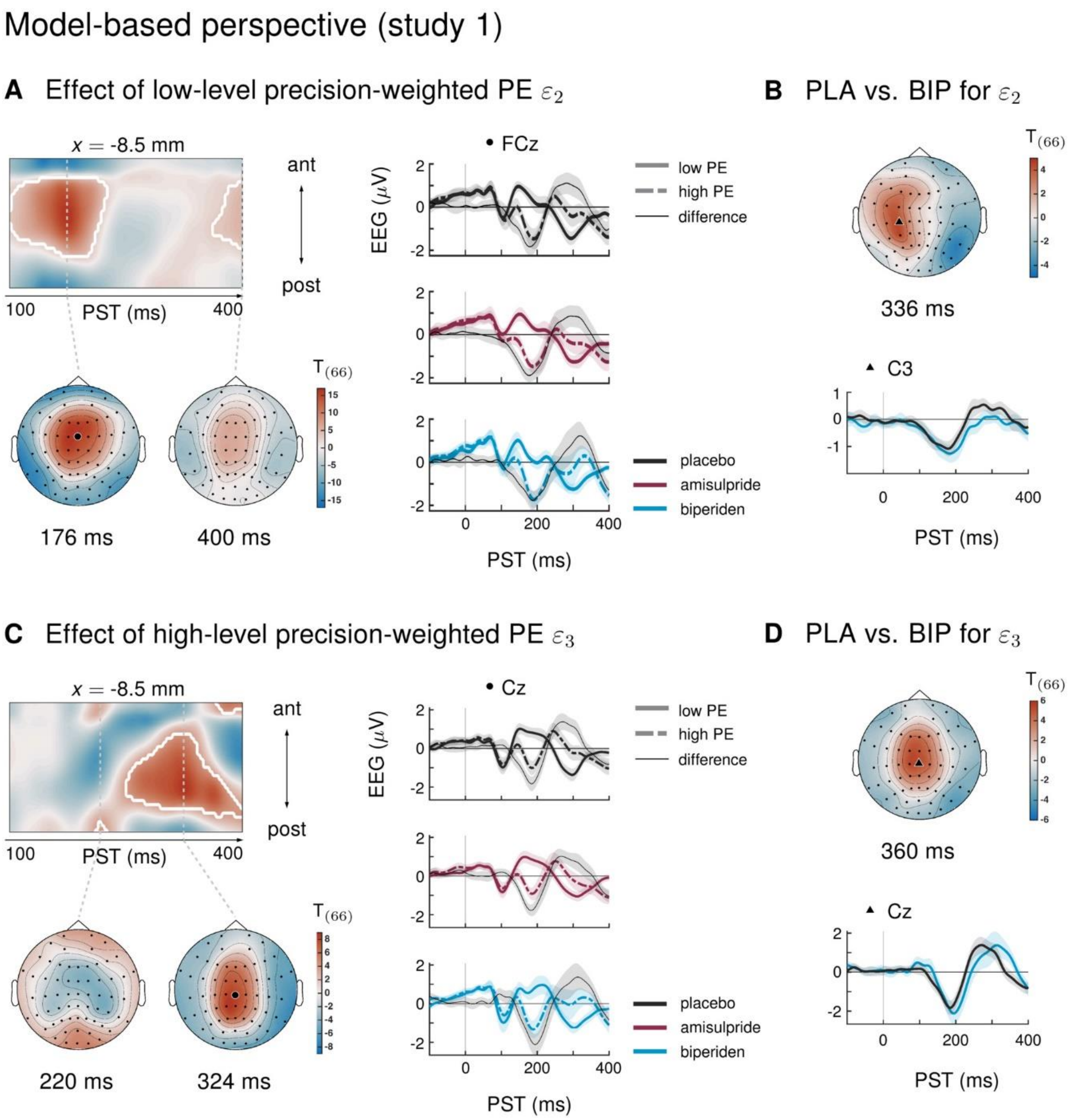


Figure S4: Results of the model-based single-trial analysis in study 1. **A** The effect of the lower-level precision-weighted PE $\varepsilon_2$ on EEG amplitudes. Left: Regions of the time × sensor space where ERPs to tones were significantly modulated by $\varepsilon_2$. Logic of display as in Figure 2 in the main text. Right: ERPs and difference waves for selected sensors, separately for the three drug groups. **B** Pharmacological effects on $\varepsilon_2$: we found a weaker modulation of ERPs by $\varepsilon_2$ under biperiden in left central sensors. Lower plot shows the ERP difference waves (ERPs to high PE tones − ERPs to low PE tones). **C** The effect of the higher-level precision-weighted PE $\varepsilon_3$ on EEG amplitudes. **D** Biperiden shifted the late central positivity encoding the higher-level PE in time.

##### *Conclusions*

As in our previously reported analysis of single-trial MMN responses in terms of hierarchical prediction errors (Weber et al., 2020), here we find that low-level precision-weighted prediction errors about stimulus occurrences are encoded in trial-wise EEG amplitudes corresponding to classical mismatch negativity, while higher-level precision-weighted prediction errors, serving as learning signals for an estimation of environmental volatility, correlate with trial-wise EEG amplitudes in a later time window. Our model-based analysis of auditory mismatch locates the dominant pharmacological effect of biperiden in this later time window, similar to the results of our whole time × sensor space analysis with an increase in trial numbers per condition (section 2.4 Alternative pre-processing and trial definition of the SM). Interestingly, under biperiden compared to placebo, we find a decrease in the correlation of EEG amplitudes with low-level PEs, but an increase in correlation with high-level (volatility) PEs, reminiscent of the relatively strong interaction effect mismatch*stability under biperiden reported in the main text.

However, when relating the results of our model-based EEG analysis to the more conventional averaging approach we followed in the main text, it is important to note that there is no reason to expect the effects of $\varepsilon_2$ and $\varepsilon_3$ to simply map onto the effects of mismatch and stability (or volatility), respectively. Specifically, the HGF is a model which already takes into account the effects of environmental volatility on belief updates by scaling the current learning rate (i.e., the precision-weight on the PE). Thus, while we did hypothesize $\varepsilon_2$ to capture the classical mismatch negativity component of the auditory ERPs, the $\varepsilon_2$ regressor itself also already scales with current volatility estimates on a trial-by-trial basis, i.e., it includes a fine-grained version of what we are trying to capture coarsely in the stability and the interaction effect (mismatch*stability) in the conventional approach.

Moreover, the higher-level PE $\varepsilon_3$ does not correspond to the current level of (estimated) volatility – this is captured in the model's variable $\mu_3$ which quantifies an individual's estimate of volatility. Instead, $\varepsilon_3$ is a belief *update* signal, which quantifies the amount to which the estimate of current speed of change in the environment (volatility) should be adjusted based on how far off the agent's belief about the statistical laws driving the stimulus occurrences was.

Importantly, while this quantity is also expected to scale with the more slowly changing volatility beliefs, it is still a function of the trial-by-trial input (e.g., shows opposite signs in response to expected tones (standards) and unexpected tones (deviants)). Because of this, it is also expected to capture ERP components which differ between standard and deviant trials – i.e., classical mismatch components, which is exactly what we find. In general, ERPs, due to their transient nature, have been hypothesized to mainly reflect such belief updates. In the specific case of auditory oddball paradigms, belief updates are believed to take place on multiple levels of a processing hierarchy. It is thus an interesting finding – and very consistent with our previous report in a different data set (Weber et al., 2020) – that later components of the auditory ERP (such as the P300) seem to reflect belief updates on a higher level, such as volatility estimation.

| **Study 1: Model** | cluster | $x[mm]$ | $y[mm]$ | $z[ms]$ | $t_{66}$ | $Z_{\equiv}$ | $p_{FWE}$ | $k_E$ | $tw_{sig}$ [ms] |
|---|---|---|---|---|---|---|---|---|---|
| Low-level PE: pos. | 1 | -60 | -57 | 176 | 14.19 | Inf | 0.000 | 5583 | 108 - 228 |
| | | -17 | 72 | 144 | 12.43 | Inf | 0.000 | | |
| | | -8 | 72 | 144 | 12.41 | Inf | 0.000 | | |
| | 2 | 47 | -36 | 360 | 6.53 | 5.71 | 0.000 | 1353 | 284 - 400 |
| | | 42 | -41 | 336 | 6.51 | 5.70 | 0.000 | | |
| | | 47 | -41 | 400 | 5.71 | 5.13 | 0.001 | | |
| | 3 | -55 | -25 | 400 | 6.06 | 5.38 | 0.000 | 423 | 360 - 400 |
| | 4 | 68 | 18 | 192 | 4.79 | 4.42 | 0.013 | 4 | 184 - 196 |
| | 5 | -42 | -30 | 312 | 4.58 | 4.25 | 0.025 | 12 | 312 - 316 |
| Low-level PE: neg. | 1 | -8 | -3 | 176 | 16.40 | Inf | 0.000 | 7361 | 100 - 232 |
| | | 4 | 29 | 168 | 15.84 | Inf | 0.000 | | |
| | 2 | -13 | -14 | 400 | 6.51 | 5.70 | 0.000 | 663 | 364 - 400 |
| Low-level PE: PLA > BIP | 1 | -26 | -30 | 328 | 4.46 | 4.15 | 0.036 | 8 | 328 - 332 |
| Low-level PE: BIP > PLA | 1 | 42 | -62 | 336 | 4.91 | 4.51 | 0.009 | 47 | 328 - 344 |
| | 2 | 38 | -62 | 392 | 4.43 | 4.13 | 0.040 | 10 | 388 - 396 |
| High-level PE: pos. | 1 | -8 | -36 | 324 | 8.48 | 6.95 | 0.000 | 3679 | 248 - 400 |
| | | -13 | -46 | 364 | 7.77 | 6.52 | 0.000 | | |
| | | -26 | -68 | 400 | 5.60 | 5.04 | 0.001 | | |
| | 2 | 4 | 72 | 392 | 6.53 | 5.71 | 0.000 | 208 | 364 - 400 |
| | 3 | 8 | -95 | 220 | 5.21 | 4.75 | 0.003 | 79 | 212 - 228 |
| | 4 | 4 | 61 | 212 | 4.46 | 4.15 | 0.036 | 11 | 208 - 212 |
| | | 21 | 61 | 212 | 4.41 | 4.11 | 0.042 | | |
| High-level PE: neg. | 1 | 64 | -52 | 328 | 6.79 | 5.89 | 0.000 | 2483 | 272 - 400 |
| | | 34 | 13 | 380 | 6.10 | 5.41 | 0.000 | | |
| | | 60 | -36 | 284 | 6.06 | 5.38 | 0.000 | | |
| | 2 | -55 | -46 | 308 | 4.85 | 4.47 | 0.011 | 165 | 296 - 320 |
| | 3 | 0 | 50 | 276 | 4.77 | 4.40 | 0.014 | 20 | 260 - 280 |
| | 4 | 30 | -30 | 216 | 4.76 | 4.40 | 0.014 | 97 | 208 - 224 |
| | | 4 | 2 | 216 | 4.59 | 4.26 | 0.024 | | |
| | 5 | 13 | -95 | 320 | 4.49 | 4.18 | 0.033 | 4 | 316 - 324 |
| | 6 | -21 | 40 | 256 | 4.41 | 4.11 | 0.042 | 2 | 252 - 256 |
| High-level PE: BIP > PLA | 1 | -8 | -14 | 360 | 5.35 | 4.86 | 0.002 | 327 | 336 - 376 |

Table S12 Results of the model-based single-trial analysis in study 1. Columns are organized as in Table 1 of the main text.

In summary, our model-based analysis provides a complementary perspective on auditory mismatch signals in our task, but agrees with the results of the conventional categorical standard/deviant approach reported in the main text in the following important points:

1. Biperiden affects auditory mismatch signals, while we find no evidence for a dopaminergic modulation of MMN by amisulpride or levodopa.

2. Biperiden does not only affect mismatch detection itself, but also the higher-level learning about the volatility of the environment – indicated by a stronger interaction effect mismatch*stability in the biperiden group (main text), as well as a significant effect of biperiden on the representation of the higher-level PE $\varepsilon_3$ which serves to update volatility beliefs.

3. Galantamine, at the dose given here, did not affect mismatch processing (or its scaling by environmental stability) in our paradigm.

#### Model-based perspective (study 2)

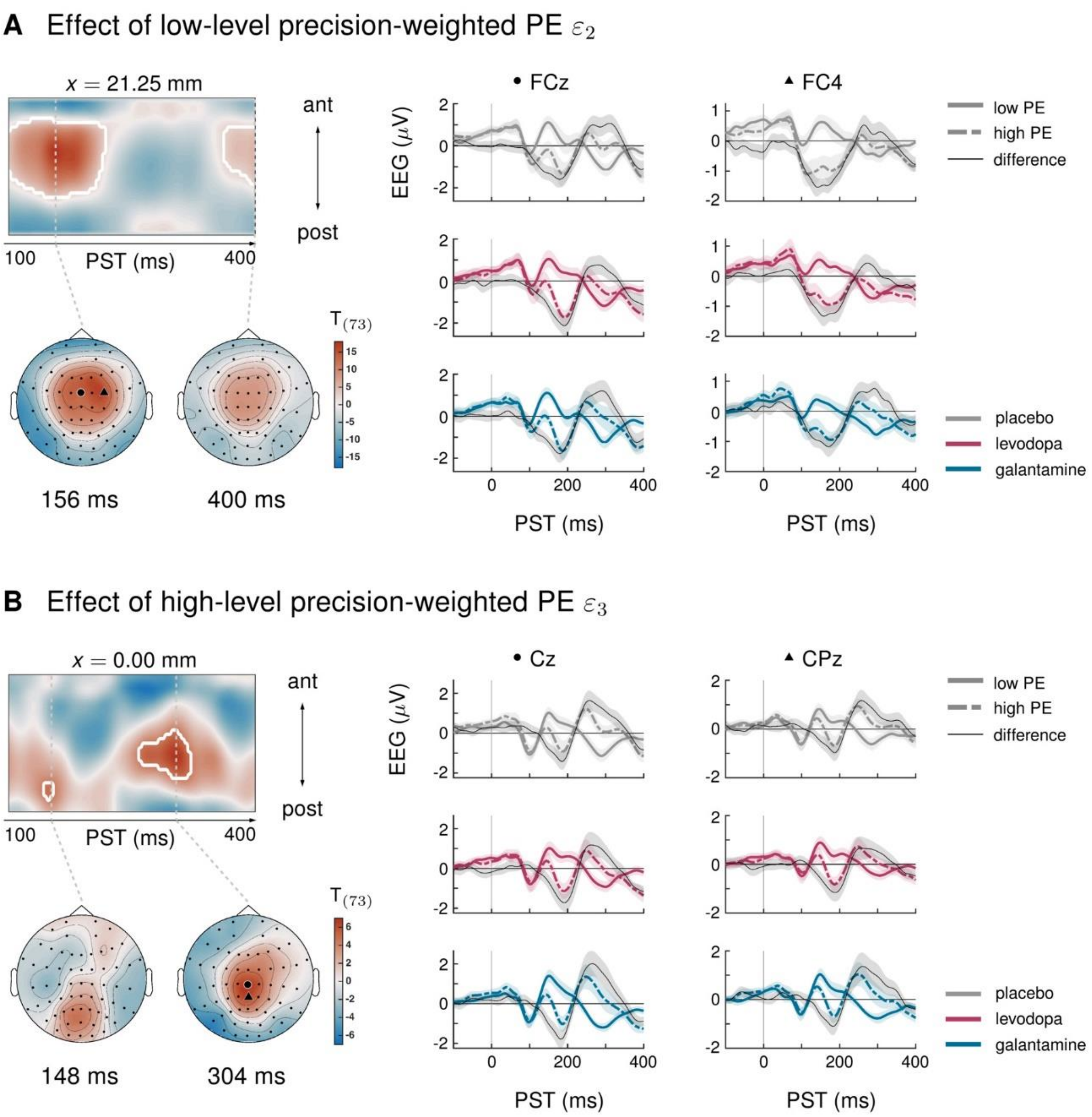


Figure S5: Results of the model-based single-trial analysis in study 2. **A** The effect of the lower-level precision-weighted PE $\varepsilon_2$ on EEG amplitudes. Left: Regions of the time × sensor space where ERPs to tones were significantly modulated by $\varepsilon_2$. Logic of display as in Figure 2 in the main text. Right: ERPs and difference waves for selected sensors, separately for the three drug groups. **B** The effect of the higher-level precision-weighted PE $\varepsilon_3$ on EEG amplitudes. There were no significant differences between drug groups in either prediction error.

| **Study 2: Model** | cluster | $x[mm]$ | $y[mm]$ | $z[ms]$ | $t_{73}$ | $Z_{\equiv}$ | $p_{FWE}$ | $k_E$ | $tw_{sig}$ [ms] |
|---|---|---|---|---|---|---|---|---|---|
| Low-level PE: pos. | 1 | -47 | -68 | 168 | 14.03 | Inf | 0.000 | 6196 | 100 - 224 |
| | | 0 | 72 | 164 | 13.51 | Inf | 0.000 | | |
| | | 47 | -73 | 192 | 12.61 | Inf | 0.000 | | |
| | 2 | -34 | -68 | 400 | 7.21 | 6.24 | 0.000 | 432 | 368 - 400 |
| | 3 | -17 | -25 | 256 | 6.85 | 6.00 | 0.000 | 1386 | 240 - 300 |
| | 4 | 0 | 72 | 400 | 6.51 | 5.76 | 0.000 | 225 | 364 - 400 |
| | 5 | 38 | -73 | 400 | 6.31 | 5.62 | 0.000 | 147 | 384 - 400 |
| | 6 | 68 | 18 | 176 | 4.51 | 4.22 | 0.032 | 5 | 168 - 184 |
| Low-level PE: neg. | 1 | 21 | 8 | 156 | 17.50 | Inf | 0.000 | 8067 | 100 - 220 |
| | | 13 | 8 | 176 | 17.07 | Inf | 0.000 | | |
| | | 34 | -3 | 124 | 11.88 | Inf | 0.000 | | |
| | 2 | -8 | -14 | 400 | 9.01 | 7.37 | 0.000 | 1582 | 360 - 400 |
| | 3 | -42 | -73 | 264 | 5.44 | 4.97 | 0.002 | 221 | 252 - 296 |
| | | -26 | -89 | 264 | 5.29 | 4.85 | 0.003 | | |
| | | -60 | -57 | 268 | 5.27 | 4.84 | 0.003 | | |
| | 4 | 42 | 50 | 324 | 5.34 | 4.89 | 0.002 | 74 | 296 - 332 |
| | | 34 | 61 | 300 | 4.74 | 4.41 | 0.016 | | |
| | | 60 | 29 | 324 | 4.68 | 4.36 | 0.019 | | |
| | 5 | 64 | -62 | 284 | 5.01 | 4.63 | 0.007 | 19 | 280 - 292 |
| | 6 | 34 | 61 | 264 | 4.40 | 4.13 | 0.044 | 2 | 264 - 264 |
| High-level PE: pos. | 1 | 0 | -25 | 304 | 6.57 | 5.80 | 0.000 | 619 | 260 - 320 |
| | | 0 | -25 | 272 | 5.38 | 4.92 | 0.002 | | |
| | 2 | -4 | -68 | 148 | 4.90 | 4.54 | 0.010 | 41 | 140 - 156 |
| High-level PE: neg. | 1 | -42 | -73 | 304 | 5.66 | 5.13 | 0.001 | 111 | 288 - 316 |
| | 2 | -4 | 56 | 268 | 5.26 | 4.82 | 0.003 | 121 | 248 - 296 |
| | 3 | 42 | -78 | 104 | 4.52 | 4.23 | 0.034 | 6 | 104 - 108 |
| | 4 | -60 | -57 | 308 | 4.48 | 4.20 | 0.038 | 2 | 304 - 308 |

Table S13 Results of the model-based single-trial analysis in study 2. Columns are organized as in Table 1 of the main text.
